# CALIPERS: Cell cycle-aware live imaging for phenotyping experiments and regeneration studies

**DOI:** 10.1101/2024.12.19.629259

**Authors:** Moises Di Sante, Melissa Pezzotti, Julius Zimmermann, Alessandro Enrico, Joran Deschamps, Elisa Balmas, Silvia Becca, Samantha Solito, Alessandro Reali, Alessandro Bertero, Florian Jug, Francesco S. Pasqualini

## Abstract

Cell cycle progression, migration, and proliferation shape development and regeneration, but simultaneous live-cell imaging remains challenging as conventional fluorescent cell cycle indicators (FUCCI) monopolize the green and red channels used by most structural and functional biosensors. To overcome this, we integrated a spectrally re-engineered FUCCI variant, open-source analysis software, and four-color human stem cell reporter lines into CALIPERS: a method for Cell-cycle-Aware Live-cell Imaging in Phenotyping and Regeneration Studies.

## Main text

The convergence of -omics and imaging techniques enabled advanced assessments of cellular phenotypes in basic science^1,2^, drug testing^3^, and regenerative medicine^4^. Further, reference human induced pluripotent stem cells (hiPSCs) and robust protocols for multilineage differentiation (e.g., cardiac muscle cells) strengthened reproducibility^5^ and extended phenotyping efforts to organoids^6,7^ and organs-on-chips^8,9^. However, the cell cycle (CC) can confound these studies because gene expression, morphology, and behavior change as cells progress from initial growth (G1 phase) to DNA replication (S), preparation for mitosis (G2), and the next cell division (M)^10^. This is well addressed in molecular phenotyping, as most -omics efforts are CC-aware thanks to the simultaneous measurement of CC genes/proteins^11^. However, only structural phenotyping of chemically fixed samples can be made CC-aware via specific markers^12^. Functional phenotyping requires live-cell imaging^13,14^, where we cannot simultaneously assess cell structure, function, and CC progression using standard fluorescence microscopes because green and red fluorescent proteins (GFP, RFP) power both Fluorescence Ubiquitin Cell Cycle Indicators (FUCCI)^10^, and most phenotypic sensors^15^. Here, we introduce an integrated framework that combines a customized FUCCI sensor (FUCCIplex), dedicated image-analysis software (FUCCIphase), and versatile genetic engineering strategies in human epithelial cells (HaCaT, Fig. 1) and hiPSCs (Fig. 2). We call this method-in-a-cell-line CALIPERS for CC-Aware Live-cell Imaging for Phenotyping Experiments and Regeneration Studies.

**Figure 1:**
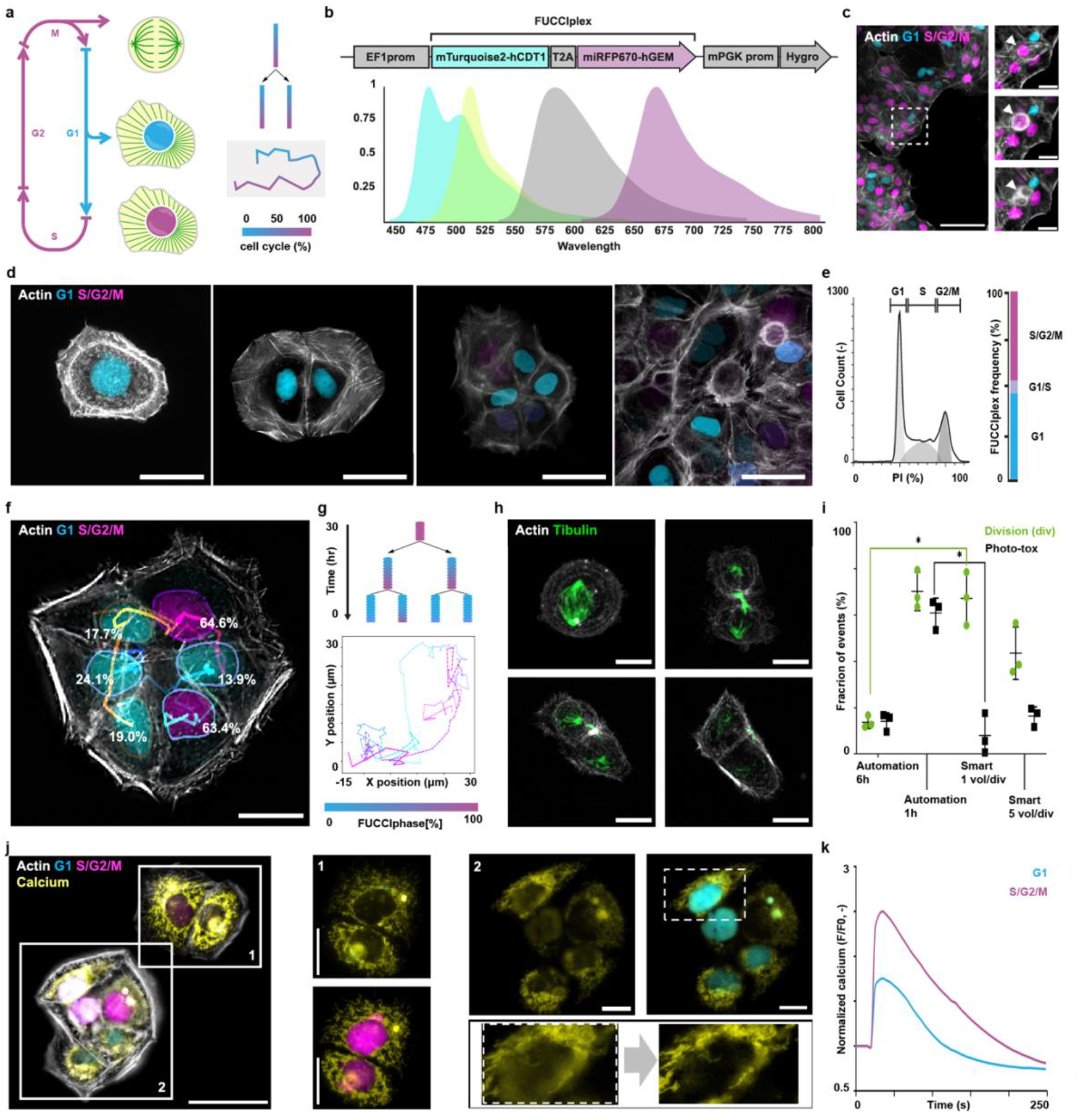
Demonstrating CALIPERS in HaCaT cells. **a)** Illustration of expected readouts from imaging the cell cycle (CC) phase together with structural (grey, actin; green, tubulin) and functional (yellow, calcium) sensors integrated in cell hierarchy and migration analyses. **b)** A schematic of the FUCCIplex construct and the multiplexable emission spectra of the fluorescent proteins used in this study. **c)** Live imaging of HaCaT cells expressing FUCCIplex (cyan/magenta) and RFP-LifeAct (grey). Insets show a cell during division. **d-e)** CC phase occupancy based on multiple static images (**d**), flow cytometry of WT HaCaT stained with PI (**e**, left) or FUCCIplex-expressing HaCaT (**e**, right). **f-g)** Example of automated segmentation and tracking (**f**) to perform a phase-locked cell hierarchy and migration analysis (**g**). **h-i)** Representative smart microscopy acquisition of mitotic spindle dynamics in FUCCIplex-LifeAct HaCaT cells with GFP-tagged β-tubulin (**h**) and statistical comparison of smart microscopy spindle yield and phototoxicity versus automatic confocal acquisitions every one or six hours (**i**). Three independent experiments, results as mean ± SD, * implies p<0.05. **j-k)** A representative live cell imaging frame (**j**) and calcium transient analysis (**k**) of HaCaT cells loaded with a calcium-sensitive dye (FLUO-4, yellow), showing no iRFP bleed-through in S/G2/M cells (panel 1), and CFP bleed-through removal in G1 cells (panel 2). Scale bars: 100 µm (a), 15 µm (b), 25 µm (elsewhere).

**Figure 2:**
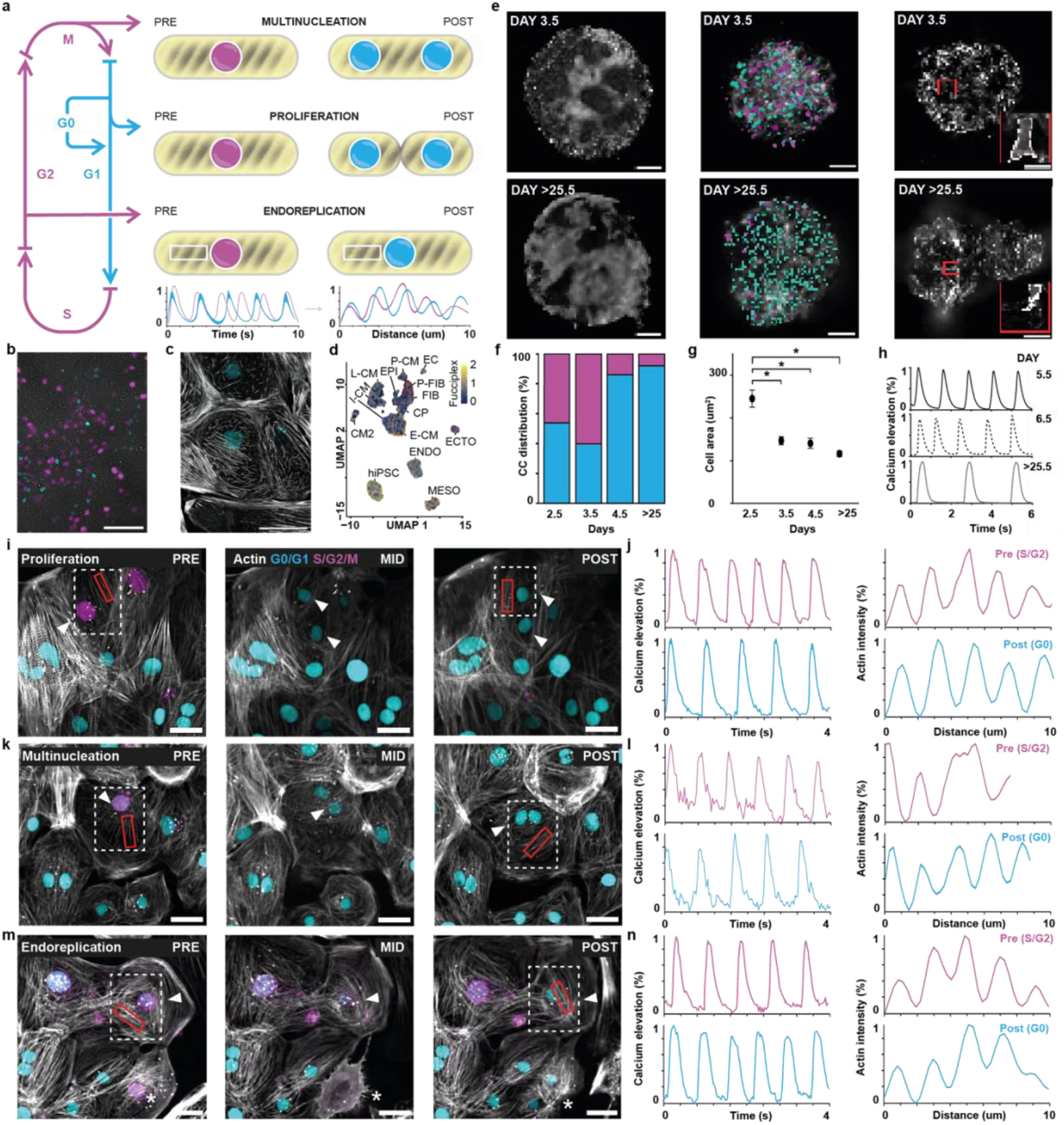
Deploying CALIPERS in hiPSCs. **a)** Illustration of expected readouts from non-canonical CC events with structural (grey, actin) and functional (yellow, calcium) phenotypic changes in hiPSC-CMs. **b-c)** hiPSC colony (**b**) and terminally differentiated (G0=constant CFP) hiPSC-CMs (**c**) expressing FUCCIplex from the EF1α and cTnT promoters, respectively. **d)** UMAP representation of single-cell RNA sequencing showing how cell types differentiated from the same hiPSC clone exhibit heterogeneous FUCCIplex expression under the EF1α promoter. Cell types include tri-lineage differentiated endoderm (ENDO), mesoderm (MESO), ectoderm (ECTO), cardioid-differentiated endothelial cells (EC), epicardial cells (EPI), proliferating fibroblasts (P-FIB), fibroblasts (FIB), cardiac progenitors (CP), early (E-CM), proliferating (P-CM), intermediate (I-CM), late (L-CM), and conductive cardiomyocytes (CM2). **e-h)** Representative light-sheet and confocal imaging of cardiac organoids obtained from genome editing CAG-FUCCIplex in the safe harbor site hRosa26 (**e**), together with CC (**f**), structural (**g**), and functional (**h**) quantifications. **i-n)** Representative images and structural and functional analyses of hiPSC-CMs re-entering the CC (iRFP-expressing nuclei) for proliferation (**i-j**), multinucleation (**k-l**), or endoreplication (**m-n**). Dashed squares and arrowheads indicate a cell pre and post event in all conditions; the asterisk indicates an additional proliferation event in the endoreplication panels. Scale bars: 100 µm (b, d), 25 µm (elsewhere).

To create FUCCIplex, we replaced the GFP/RFP pair in fast-FUCCI^16^ with miRFP670 (iRFP) and mTurquoise2 (CFP). Thus, FUCCIplex avoids toxic UV excitations, unlike a previous attempt^17^; works with standard filter sets, unlike TMI^18^; and frees the green/red channels used by the majority of biosensors, unlike Fucci4^19^. Practically, the nucleus of FUCCIplex cells contains CFP in G1, both CFP and iRFP in the G1-S transition, and only iRFP during the S/G2/M phases (Fig. 1a-b). To showcase CALIPERS, we co-expressed FUCCIplex and the actin-binding peptide RFP-LifeAct^20^ in HaCaT cells and used live-cell fluorescence imaging over 40 hours to track CC phase duration (Fig. 1c and Supplemental Video 1-2). We confirmed that HaCaT cells spent ∼40% of their time in G1 and the rest in S/G2/M, in good agreement with the occupancies measured both with static FUCCIplex images and flow cytometry experiments with a DNA marker (Fig. 1d-e and Extended Fig. 1). Further, we developed an open-source plug-in, FUCCIphase, that calculates CC percent completion based on nuclear intensities, enabling CC-aware morphology and motility analyses (Fig. 1f-g, Supplemental Video 3, and Extended Fig. 2).

To build four-color CALIPERS reporter lines while avoiding UV-induced CC photodamage^17^, we had to accept a limited spectral overlap between CFP and GFP (Fig. 1b). To minimize artifacts, we deployed two complementary strategies. First, we leveraged the lack of bleed-through in S/G2/M for late CC events, e.g., mitotic spindle formation^21^. We fused GFP to β-tubulin to enable a smart microscopy routine that maximizes spindle acquisitions and minimizes phototoxicity by switching from 2D widefield to 3D confocal imaging when FUCCIphase > 90%. Over 24 hours, we observed phototoxicity comparable to that of automatically acquiring a volume every six hours, but spindle yield on par with imaging a volume every hour. Thanks to the reduced phototoxicity, we acquired up to five volumes per division (Fig. 1h-i, Extended Figs. 3-4, and Supplementary Video 4). The second strategy, compatible with all CC phases, combines FUCCIplex and LifeAct with a faster GFP-based calcium sensor, as calcium time courses are orders of magnitude faster than the CC. To demonstrate this approach, we treated HaCaT cells with a GFP-based calcium-sensitive dye, imaged ATP-induced calcium releases, and automatically extracted CC-aware morphological and intensity-based features after removing the static artifact (Fig. 1j-k, Supplemental Video 5, and Supplemental Fig. 2).

Having validated CALIPERS in HaCaT cells, we next implemented it in hiPSCs to facilitate adoption across differentiated cell types. We focused on hiPSC-derived cardiomyocytes, where non-canonical CC events such as terminal differentiation (G0), endoreplication, and multinucleation complicate phenotypic interpretation (Fig. 2a)^22^. Furthermore, the changes in cell structure and function during these events are poorly understood but critical for regenerative therapies^23,24^. To address this, we generated a novel dual-reporter line expressing RFP-LifeAct and GCaMP6f to visualize contractile myofibrils and calcium cycling (Extended Fig. 5)^13,14^. Then, we established lentiviral (LV) and genome-editing delivery strategies to create a method-in-a-cell-line compatible with post-differentiation labeling^8^, or stable expression across lineages^25^.

Using LVs with constitutive (EF1α, CMV) and cardiac-specific (cTNT, hTNNT2) promoters (Extended Fig. 6), we confirmed promoter-dependent FUCCIplex expression in hiPSC (Fig. 2b) and hiPSC-CMs (Fig. 2c), with the stem cells cycling normally and hiPSC-CMs exhibiting constant CFP-positive nuclei characteristics of cells in G0^22^. While EF1α uniquely supported FUCCIplex expression in undifferentiated hiPSCs (Supplementary Video 6), single-cell RNA sequencing of hiPSC-derived cells from germ layers and cardiac organoids revealed substantial expression heterogeneity (Extended Fig. 7). Notably, the silencing in hiPSC-CMs could be leveraged to enable further microtubule multiplexing^26^ (Supplementary Video 7 and Supplementary Figure 3) and a nocodazole drug test in proliferating vs non-proliferating hiPSC-derived cell types (Extended Fig. 8, Supplementary Video 8-9).

Next, we genome-edited a stable four-color reporter line by inserting FUCCIplex into the human Rosa26 locus^25^ (Extended Fig. 9) and repeated the cardiac organoid experiments (Fig. 2e, Supplementary Videos 10-12, and Supplemental Figure 3). While all cells in the cardioid retained FUCCIplex expression, many hiPSC-CMs had exited the CC by day 4.5. By day 25, virtually all were post-mitotic (Fig. 2f). The area of hiPSC-CMs also decreased sharply after organoid formation (day 3.5–4.5), with only modest further reductions as pseudo-chambers consolidated, reflecting developmental compaction (Fig. 2g). Finally, regular calcium transients appeared by day 5.5, shortly after cell-cycle exit and structural maturation (Fig. 2h). This progression from proliferation exit to structural reorganization to functional onset tracked cardiac development in-vivo^27^. Finally, in cardiac regenerative medicine, we seek therapies that induce CC re-entry^23,24^ while integrating mature structure and function at the regeneration site^4,28^. By detecting rare iRFP-positive nuclei, we automatically identified cycling hPSC-CMs and performed CC-aware structural and functional phenotyping pre- and post-proliferation (Fig. 2i-j), multinucleation (Fig. 2k-l), and endoreplication (Fig. 2m-n, Supplemental Video 13) events^22,29^. Notably, these analyses could be extended in content and throughput (Extended Fig. 10 and Supplementary Videos 14-15), suggesting CALIPERS might screen regenerative interventions in the future^4^.

In summary, CALIPERS offers CC-aware structural and functional cellular phenotyping via three plug-and-play adoption routes. First, as a method-in-a-cell-line, the turnkey four-color WTC-11 reporter simultaneously tracks actin architecture, calcium dynamics, and CC phase. Second, FUCCIplex can be integrated onto any existing GFP/RFP reporter line via lentiviral delivery or CRISPR safe-harbor knock-in to retrofit the CC context. For example, we validated FUCCIplex in WTC-11 to ensure compatibility with the Allen Cell Collection, the field’s largest panel of GFP/RFP hiPSC reporters. Third, users may start from the blank-canvas FUCCIplex WTC-11 line and add their favorite GFP/RFP sensors to build new CC-aware models. Across all implementations, the open-source FUCCIphase pipeline provides a unified analysis backbone and remains fully compatible with legacy two-color FUCCI data. Overall, we are confident that CALIPERS will extend to live-cell imaging the same advantages that cell-cycle awareness has already brought to –omics disciplines.

## Supporting information

CALIPERS datafile 1

CALIPERS datafile 2

## Acknowledgement

FSP gratefully acknowledges funding from the European Research Council (ERC StG #852560 and the Chips Joint Undertaking (Grant Agreement No. 101140192 to FSP) and its members, including the top-up funding of Belgium, Germany, Hungary, Ireland, Italy, the Netherlands, Portugal, Romania, and Spain. For Italy, the work was funded by the Italian Ministry of Enterprises and Made in Italy (MIMIT) under CUP F13C23003260003. While the work was supported by the European Union, the views and opinions expressed are those of the author(s) only and do not necessarily reflect those of the European Union or Chips Joint Undertaking. Neither the European Union nor the granting authority can be held responsible for them.

## Extended figures

**Extended Figure 1:**
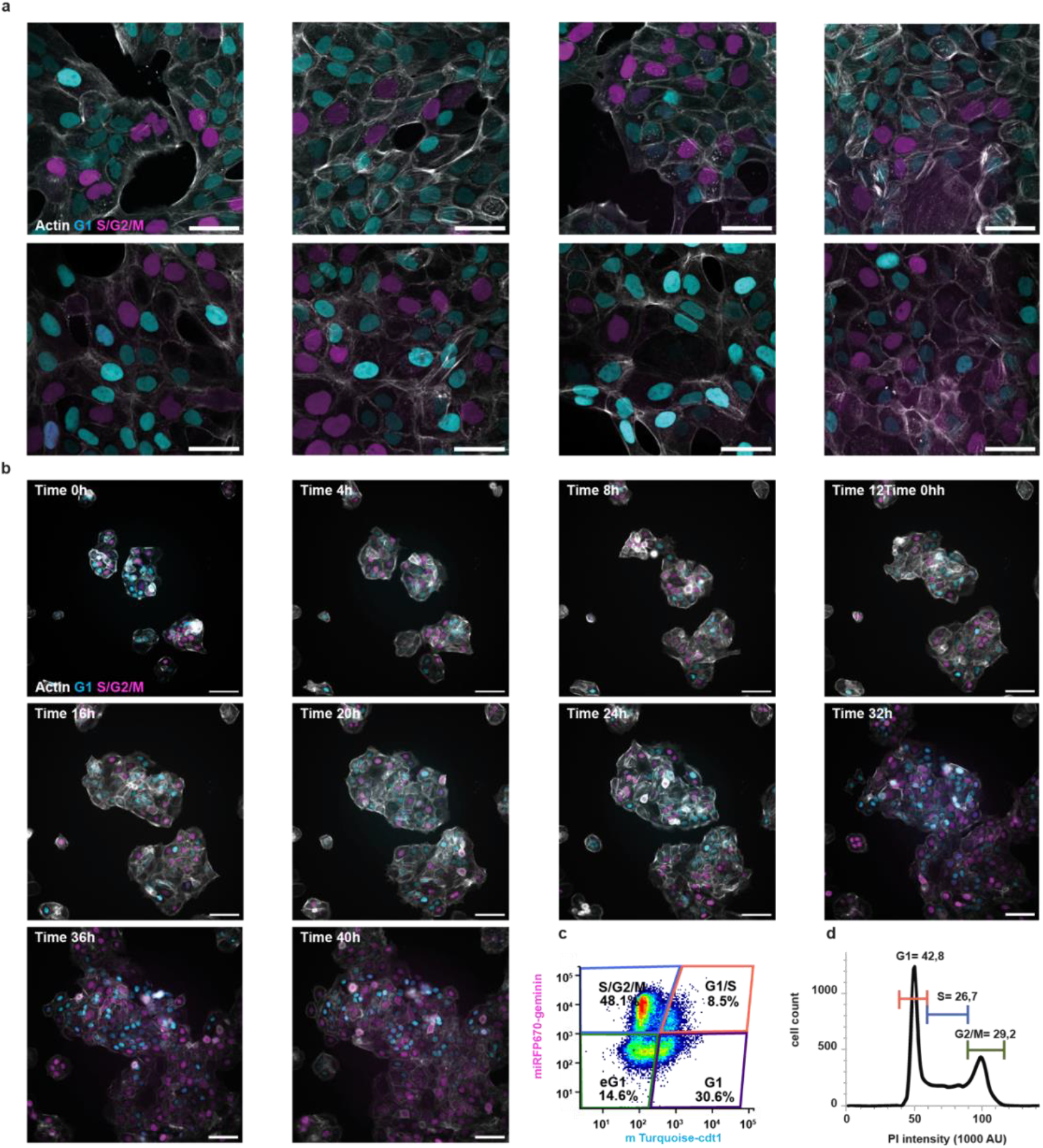
Deploying FUCCIplex in HaCaT cells. **a)** Live imaging of eight different clusters of HaCaT cells expressing FUCCIplex (cyan/magenta) and RFP-LifeAct (grey). **b)** Time-lapse live cell imaging acquisition of HaCaT cells expressing FUCCIplex and RFP-LifeAct, followed for 18 hours. **c)** CC analysis by flow cytometry based on FUCCIplex fluorescence. **d)** CC phase occupancy obtained by flow cytometry (propidium iodide staining). Data in this panel is also shown in abbreviated form in the manuscript’s Fig. 1. Details on the gating strategy are in Supplemental Fig. 1. Scale bars: 50 μm (a) and 1 μm (b).

**Extended Figure 2:**
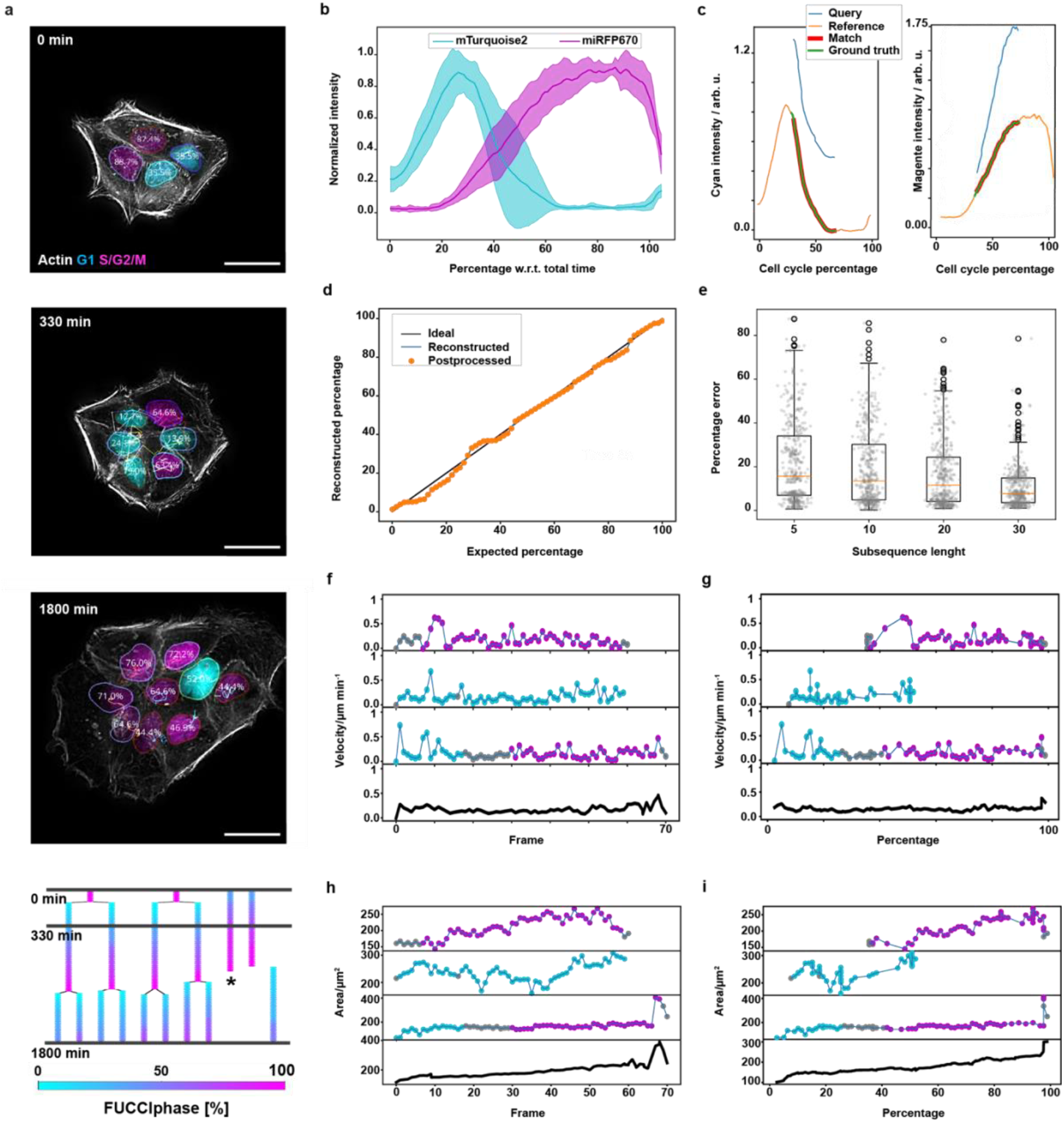
The FUCCIphase algorithm and phase-locking motility analysis. **a)** Representative frames from a proliferation video showing tracked nuclei with a CC percentage estimate at each time point determined by the FUCCIphase algorithm. At the bottom, the lineage tree from this movie is color-coded according to estimated CC percentage. A frame from this analysis is also shown in the manuscript’s Fig. 1. **b)** The reference curve for HaCaT cells is distilled from 11 tracks covering the entire CC. **c)** An example of sub-sequence matching was used to query the CC percentage from the reference curve (panel b) based on the current intensity profiles. The matched subsequence is also shown in comparison to the ground truth subsequence. **d)** Comparison of the reconstructed versus the expected percentage over a full CC, highlighting a postprocessing step to obtain percentages with nonnegative time derivatives. **e)** Percentage error per sequence depending on subsequence length: Thirty subsequences were sampled per track, and a boxplot of the CC percentage error compared to the CC percentage computed from the whole track is shown. **f)** Centroid velocity as a function of the elapsed time in frames. **g)** Centroid velocity as a function of the estimated CC percentage. **h)** Nucleus as a function of the elapsed time in frames. **i)** Nucleus area as a function of the estimated CC percentage. **j)** Scale bar: 25 μm (a).

**Extended Figure 3:**
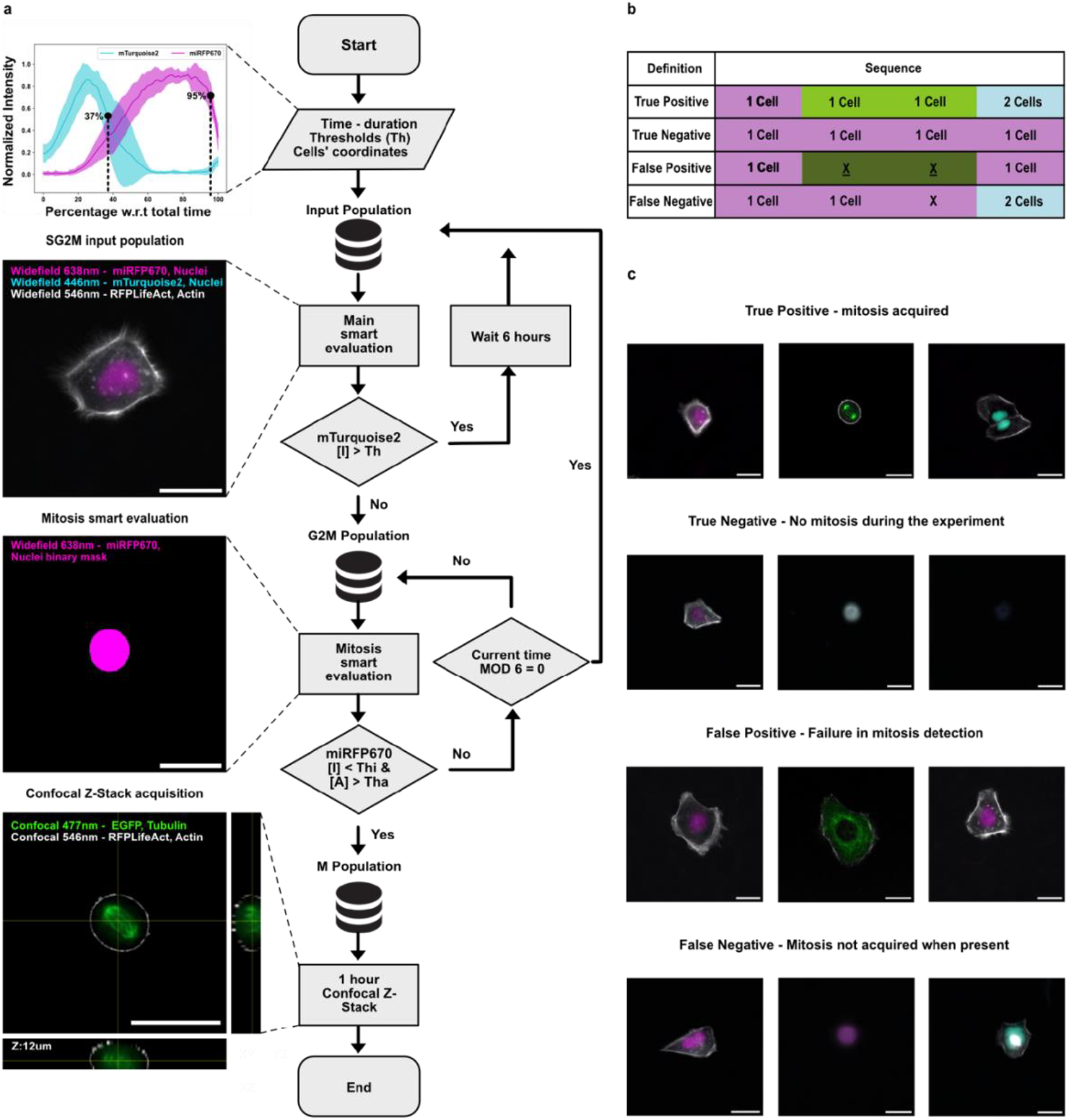
Development and validation of FUCCIsmart. **a)** Flowchart illustrating our FUCCIplex-based smart microscopy (FUCCIsmart) routine as implemented in NIS Elements JOBs Module. **b)** Table summarizing the FUCCIsmart classification method within the color-coded CC phase sequence. **c)** Representative live imaging example of FUCCIsmart classifications within HaCaT cells with GFP-tagged β-tubulin, RFP-tagged LifeAct, and FUCCIpleax. Scale bar: 25 µm.

**Extended Figure 4:**
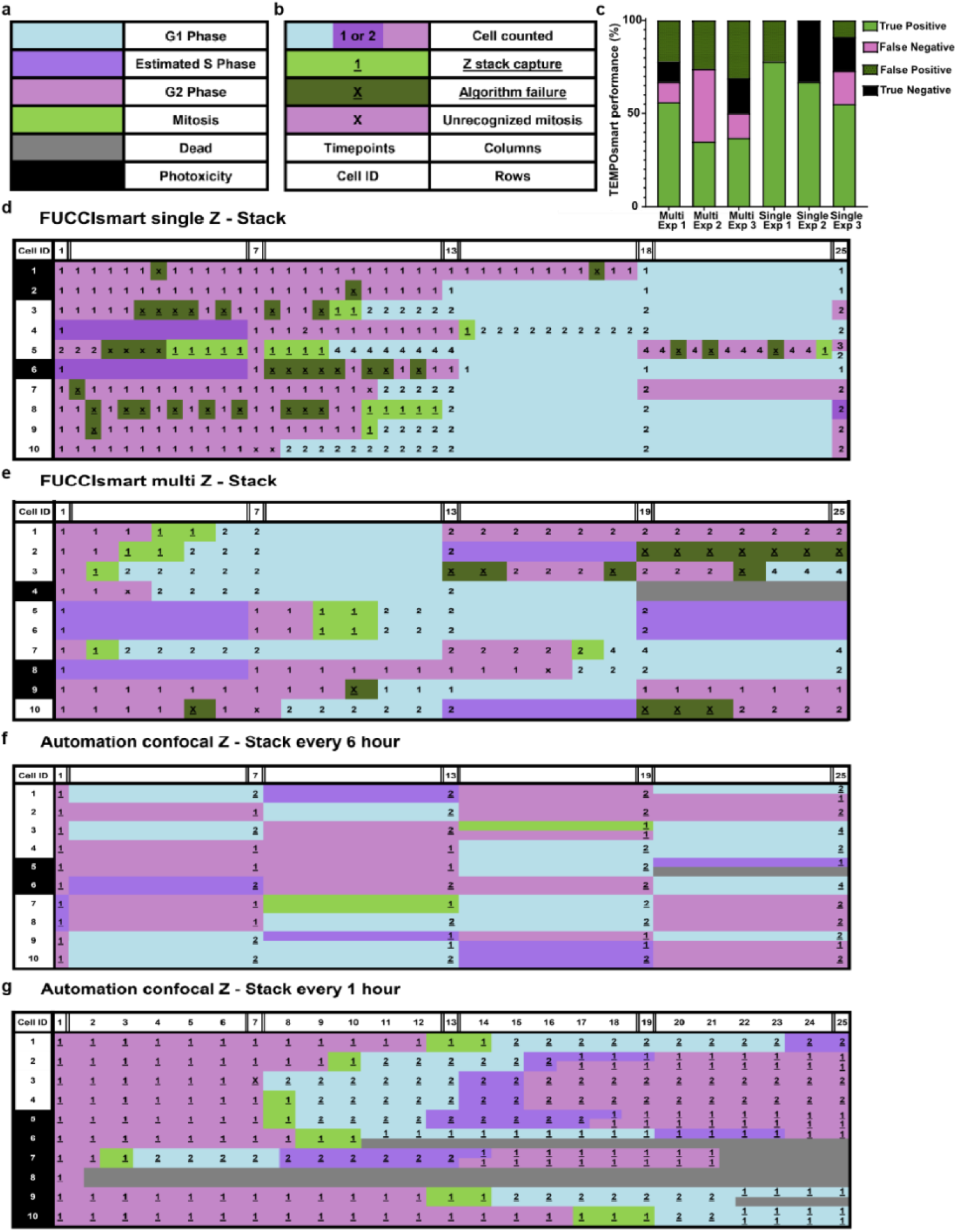
FUCCIsmart performance evaluation tables. **a-b)** Summary tables of main (**a**) and secondary (**b**) events during the confocal and smart acquisitions. **c)** The bar graph displays the percentages of True Positives (cell division was captured by 3D confocal), False Positives (3D acquisition trigged without a cell division), True Negatives (cell did not divide but 3D acquisition never triggered), and False Negatives (cell division occurred but 3D acquisition did not) for both routines (see Extended Methods in the Supplementary Information file for details on how True/False positives/negatives were assigned). **d-g)** Data acquisition sequences in FUCCIsmart routines for single (**d**) and multiple (**e**) acquisitions compared to confocal automation every six hours (**f**) and one hour (**g**). Rows represent individual cell recordings over 24 hours; columns correspond to the acquisition time points. CC phases are color-coded according to FUCCIplex labeling. The FUCCIsmart performance was evaluated across three independent experiments.

**Extended Figure 5:**
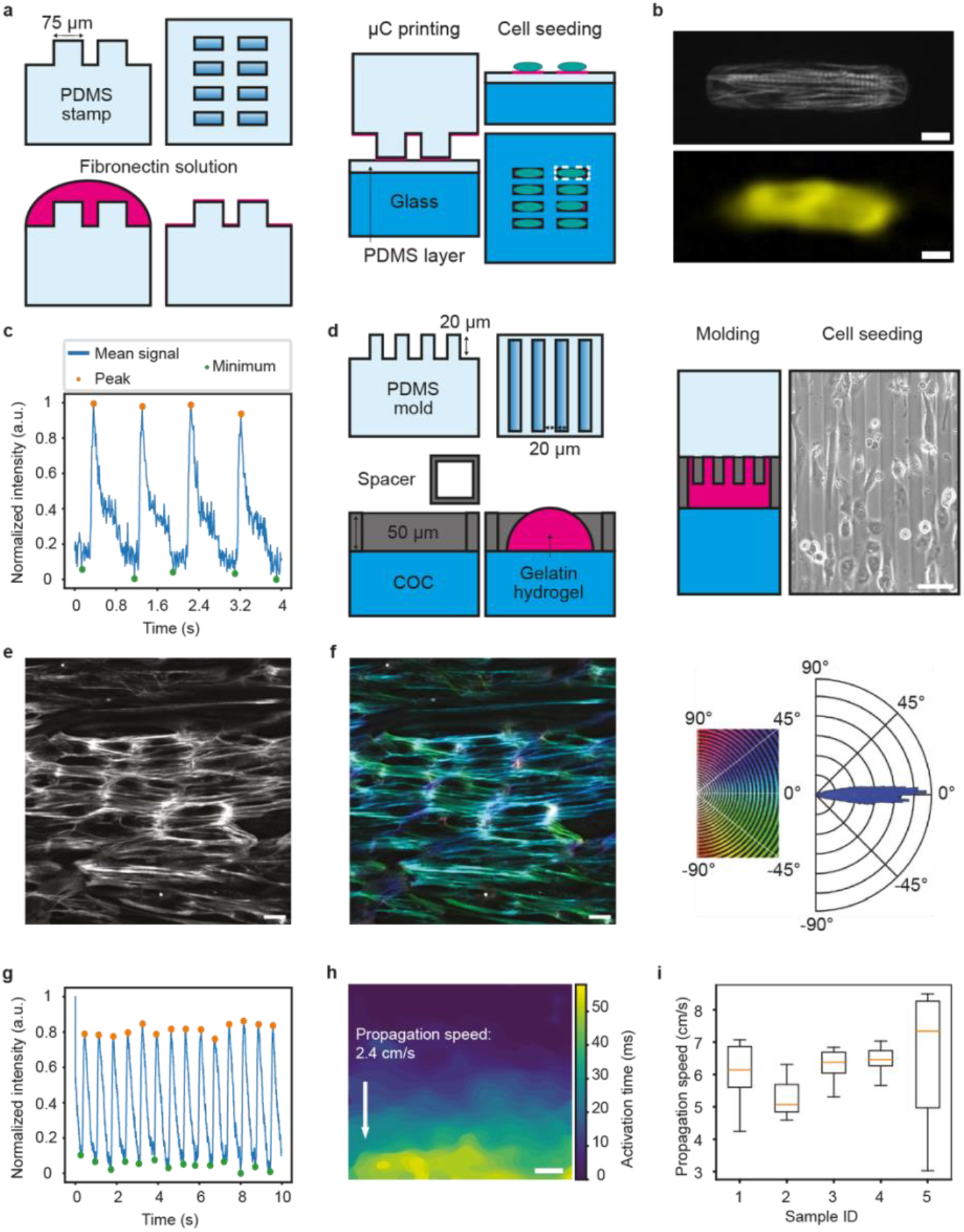
two-color reporter hiPS-derived cardiomyocytes on engineered substrates. **a)** Schematic illustration of the microcontact printing process to ink PDMS stamps featuring cell-guiding cues and transfer the resulting fibronectin islands onto glass substrates. **b)** A confocal fluorescent image of hiPSC-CMs six days after seeding shows the calcium (yellow) and F-actin (grey) signals. **c)** Representative dynamic of calcium transient signal obtained at 25 fps recording. **d)** Schematic illustration of the micro molding process to obtain gelatin hydrogels with molded line arrays for CM tissue alignment and brightfield image of hiPSC-CMs seeded on the molded gelatin gels. **e)** Confocal fluorescent image of F-actin signal. **f)** Actin alignment analysis using OrientationJ ImageJ plugin, with colormap and radar plot showing cell alignment along the molded lines. **g)** Synchronous calcium transients in aligned hiPSC-CMs obtained at 500 fps. **h)** A representative chronomap displaying the activation time for the calcium transient obtained in g. **i)** Estimation of the average propagation speed of the calcium signal in the transversal direction with respect to the tissue alignment. Scale bars: 10 μm (b), 50 μm (d), 20 μm (e, f) and 200 μm (h).

**Extended Figure 6:**
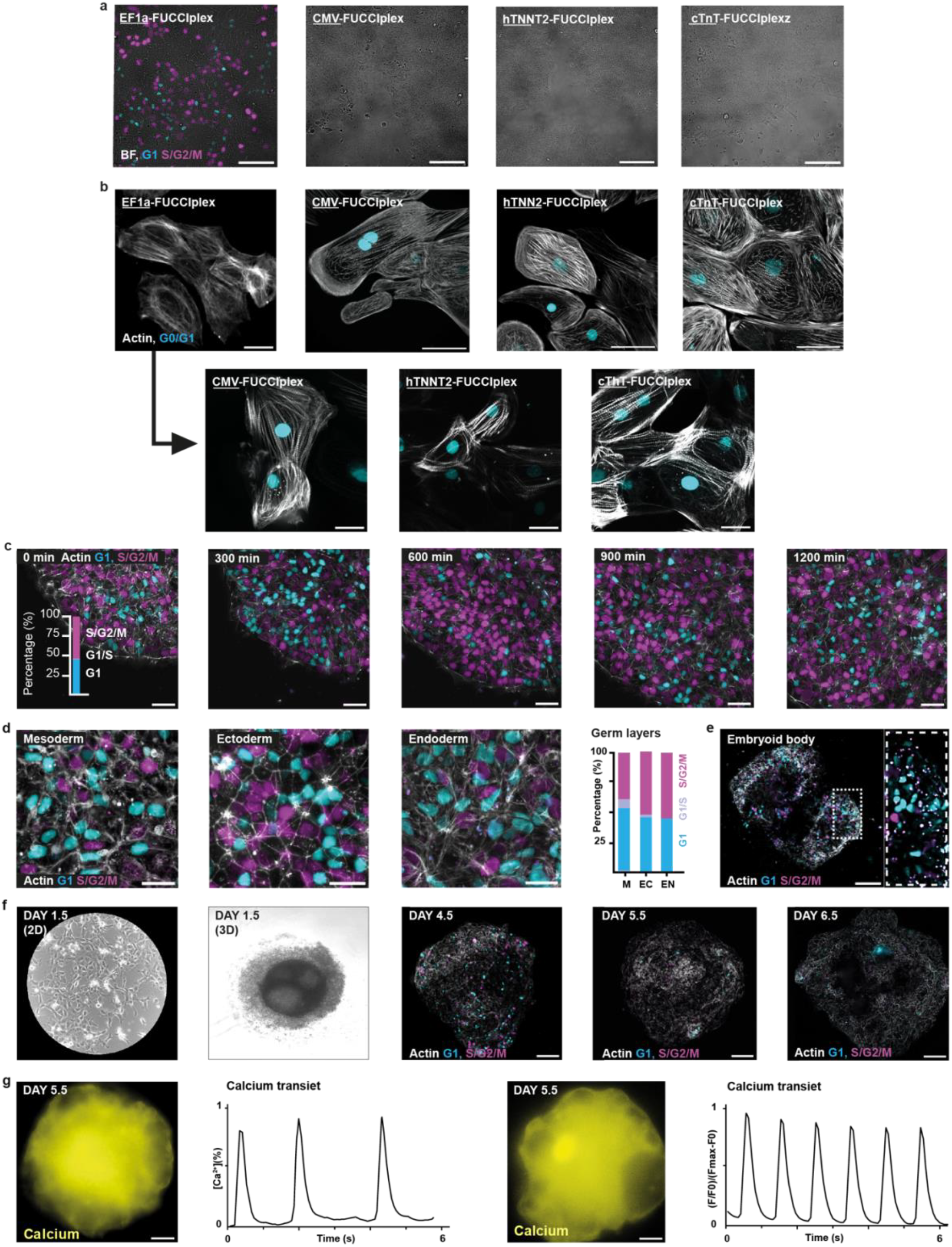
Lentiviral transduction of different FUCCIplex constructs in hiPSC-CMs. **a)** Expression of FUCCIplex in hiPSCs engineered to express the calcium and actin sensor (Extended Fig. 5) under the control of constitutive (EF1α and CMV) and cardiac-specific (hTNNT2 and cTnT) promoters **b)** Expression of FUCCIplex under the same promoters directly in hiPSC-CMs (top row) and re-expression of the same viral vectors in hiPSC-CMs differentiated from EF1α-FUCCIplex hiPSCs to show that silencing is due to the silencing of the original construct. **c)** A 20-hr, time-lapse live cell imaging acquisition of hiPSCs expressing EF1*α*-FUCCIplex (cyan/magenta) and RFP-LifeAct (grey) with quantification of how many cells are in each CC phase (inset). **d)** Live imaging of the three germ layers derived from the hiPSC clone with their CC distribution. **e)** Expression of FUCCIplex and RFP-LifeAct in embryoid bodies generated from hiPSCs (day 14). **f)** Generation of cardiac organoids derived from hiPSCs with the passage from 2D to 3D at day 2.5. Expression of FUCCIplex, GCaMP6f, and RFP-LifeAct (grey) at days 4.5, 5.5, and 6.5. **g)** Representative calcium transients are reported only for beating organoids (days 5.5 and 6.5). Scale bars: 100 µm (e-g), 50 µm (a, c), and 25 µm (b).

**Extended Figure 7:**
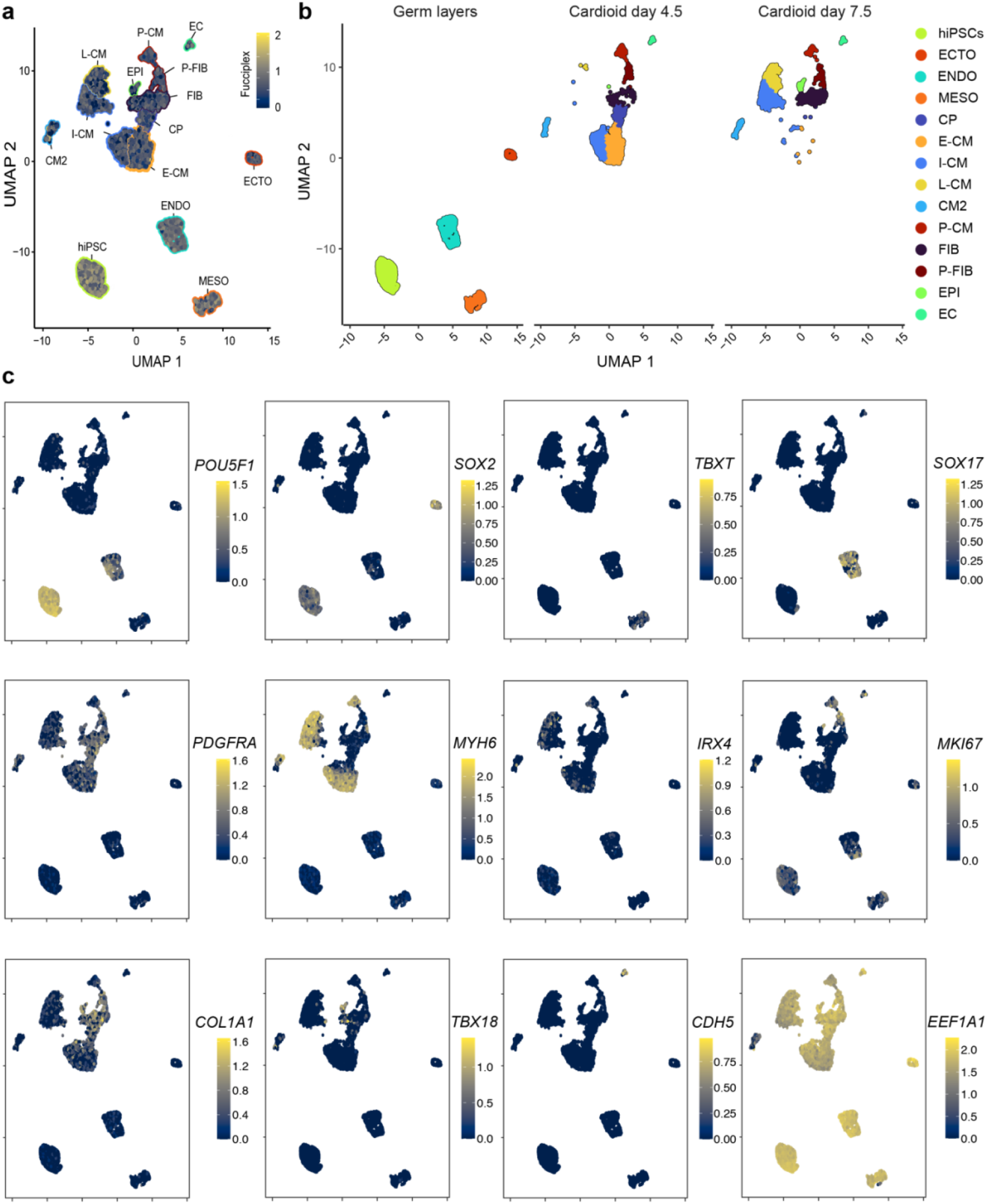
Single-cell RNAseq analysis of human cardioids. **a)** Dimensionality reduction and clustering showing FUCCIplex expression under the EF1a promoter in the main cell states and associated cell types together with an alternative UMAP representation with a breakdown of the cell states and associated cell types in the three experimental groups. **b)** Fingerprinting across hiPSC, germ layers, and cardioids after 4.5 and 7.5 days of differentiation. Cell types include tri-lineage differentiated endoderm (ENDO), mesoderm (MESO), ectoderm (ECTO), cardioid-differentiated endothelial cells (EC), epicardial cells (EPI) proliferating fibroblasts (P-FIB), fibroblasts (FIB), cardiac progenitors (CP), early (E-CM), proliferating (P-CM), intermediate (I-CM), late (L-CM), and conductive cardiomyocytes (CM2). **c)** Individual plots showing the relative expression of lineage-defining genes in the cell types represented in **a-b**.

**Extended Figure 8:**
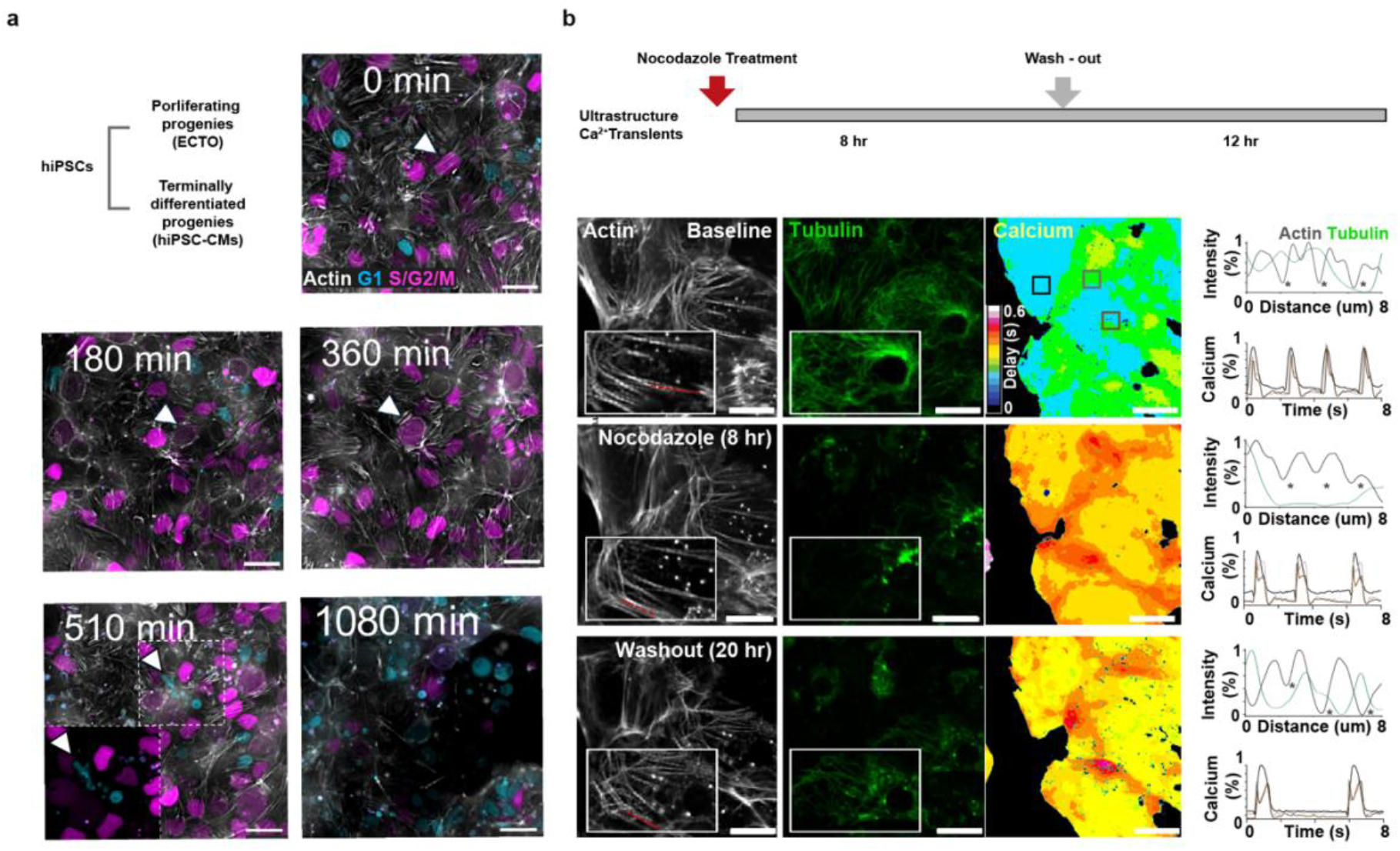
Drug testing with EF1a-FUCCIplex hiPSC-CMs. **a)** Live imaging acquisition of ectoderm cells treated with Nocodazole and imaged every 30 minutes for 18h. **b)** Structural (actin: grey, tubulin: green) and functional (calcium: yellow) analysis of nocodazole effect on EF1α-FUCCIplex hiPSC-CMs. Scale bars: 25 µm.

**Extended Figure 9:**
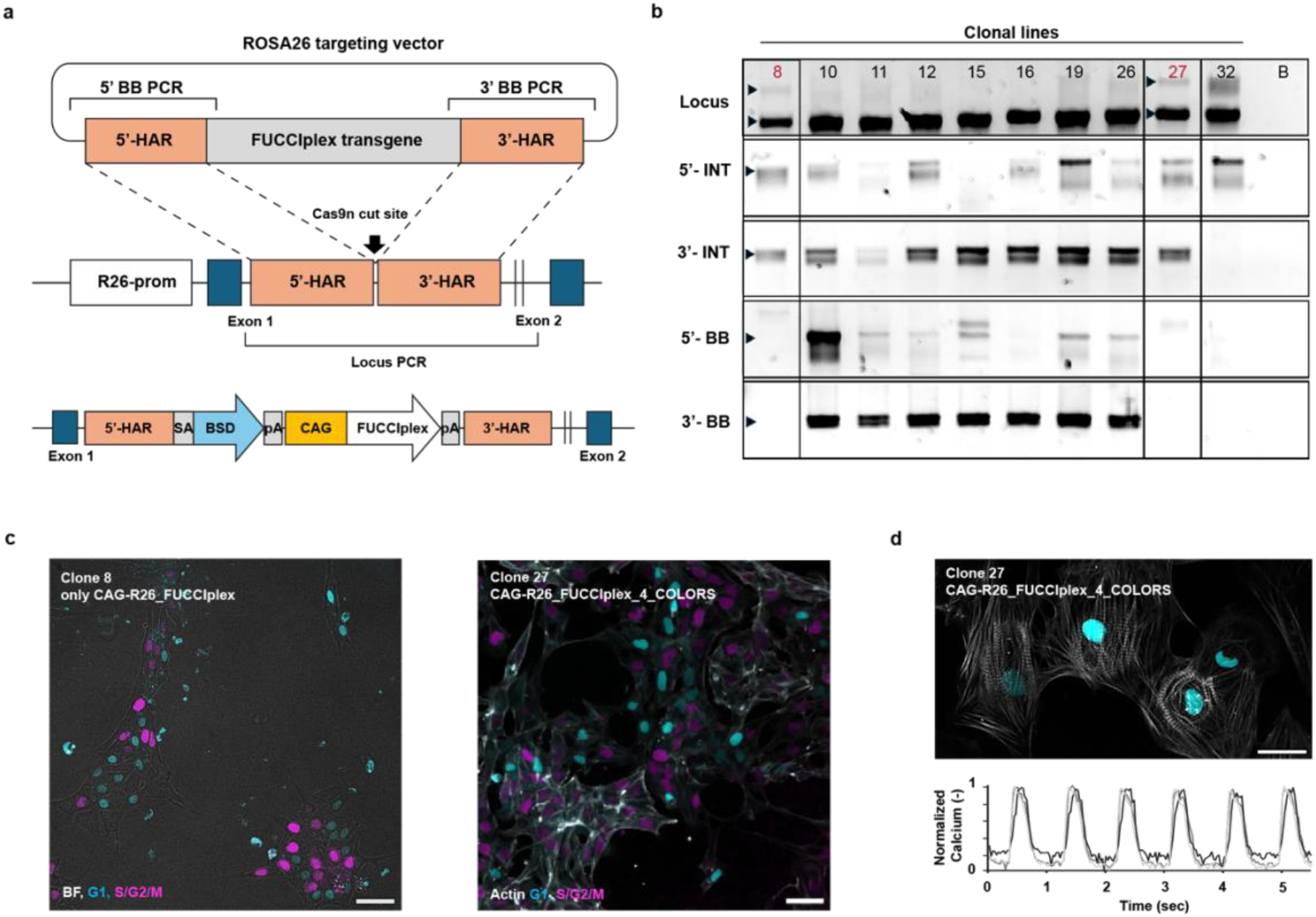
hRosa26-CAG-FUCCIplex hiPSC-CMs. **a)** Schematic diagram of the genomic insertion of the FUCCIplex construct in the ROSA26 locus. The ROSA26 targeting vector bears the FUCCIplex transgene flanked by the left and right ROSA26 homology arms (HA). Upon homology recombination, the genome-edited cells acquire Blasticidin resistance (BSD) and the FUCCIplex transgene, both under the control of a CAG promoter. **b)** PCR shows the wild-type locus, the correct 5’ and 3’ integration (INT) of the FUCCIplex construct, the 5’ and 3’ off-target backbone integration (BB), and blank (B). Clone 8 (hiPSC edited only with FUCCIplex, dual color) and clone 27 (FUCCIplex positive, GCaMP6f and LifeAct-RFP, four colors) are heterozygous and no BB insertion was detected. **c-d)** Expression of hROSA26-CAG-FUCCIplex in clone 8 (only FUCCIplex) and clone 27 (four colors) hiPSC and hiPSC-CMs. BF: bright-field. Calcium traces of three different cardiomyocytes derived from clone 27. Scale bars: 50 µm (c) and 25 µm (d).

**Extended Figure 10:**
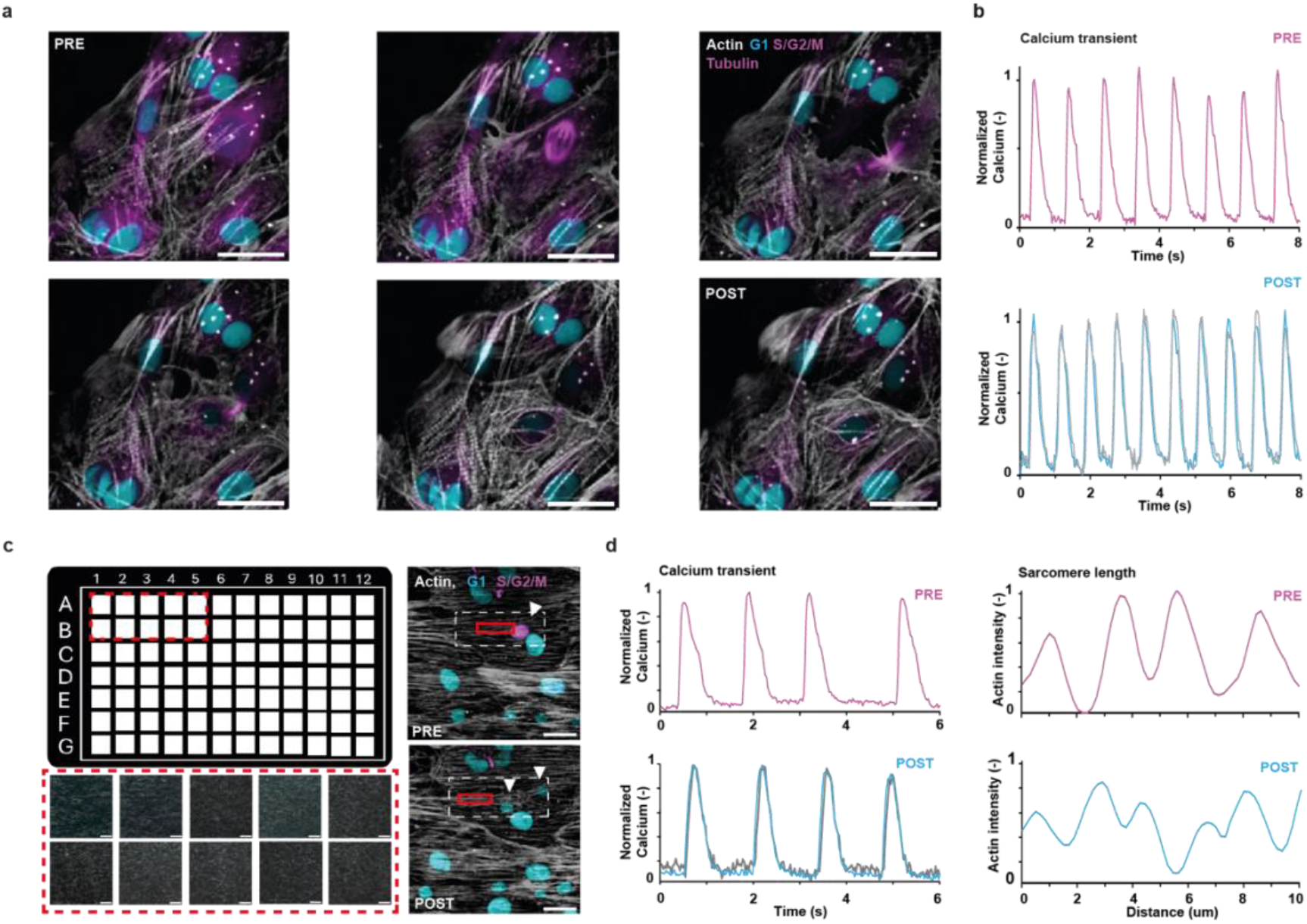
hRosa26-CAG-FUCCIplex hiPSC-CMs. **a)** Four-color FUCCIplex hiPSC-CMs were imaged after loading with a SPY650-tubulin probe. Since the iRFP intensity is dim during the M-phase in dividing hiPSC-CMs, we can clearly distinguish mitotic spindle dynamics in the nuclei (arrowhead). Calcium transients were recorded before (PRE) and after (POST) cell division. **b)** Four-color FUCCIplex hiPSC-CMs plated in a 96-well plate with a nano-grooved surface. After cardiomyocyte alignment, calcium transients and sarcomere length were quantified before (PRE) and after (POST) cell division (dashed square). Scale bars: 25 µm.

## Notes

### Competing Interest Statement

The authors have declared no competing interest.

### Summary of Updates

We strengthened the characterization of the CALIPERS hiPSCs, added a validation in cardiac organoids imaged with light-sheet microscopy, which led to a phenotyping effort of growing organoids using the four-color reporter line. We also updated the main figures and added/revised supplemental videos.

