## Supplementary material for "CALIPERS: Cell cycle-aware live imaging for phenotyping experiments and regeneration studies": Online methods

**Abstract:**

Cell cycle progression, migration, and proliferation shape development and regeneration, but simultaneous live-cell imaging remains challenging as conventional fluorescent cell cycle indicators (FUCCI) monopolize the green and red channels used by most structural and functional biosensors. To overcome this, we integrated a spectrally re-engineered FUCCI variant, open-source analysis software, and four-color human stem cell reporter lines into CALIPERS: a method for Cell-cycle-Aware Live-cell Imaging in Phenotyping and Regeneration Studies.

**Word count: 1678/1500**

*FUCCIplex plasmid engineering.* The FUCCIplex reporter was derived from pBOB-EF1-FastFUCCI-Puro by exchanging mKO2 (Cdt1 degron) with mTurquoise2 and hmAzami Green (Geminin degron) with miRFP670, thus freeing the green/red spectrum for additional sensors. The cassette is driven by the human EF1α promoter and cloned into a third-generation lentiviral backbone that carries an mPGK-Hygromycin resistance module. Promoter-specific variants (CMV, hTNNT2, cTNT) were produced by replacing EF1α with the indicated regulatory elements using standard restriction/ligation, yielding pLV[Exp]-Hygro-CMV-FUCCIplex and related constructs. All junctions were sequence-verified and VectorBuilder packaged the final vectors. FUCCIplex lentiviral vectors (pLV[Exp]-Hygro-EF1A-FUCCIplex; CMV, hTNNT2 and cTNT derivatives) will be distributed through Addgene (new constructs pending ID).

*Lentiviral production.* HEK293T packaging cells ($2.5 \times 10^6$ cells per 10 cm dish; 0.2 % gelatin-coated) were co-transfected with the FUCCIplex transfer plasmid and third-generation helper plasmids using Lipofectamine 2000. Viral supernatants were collected 48 h post-transfection, clarified, and applied to target cells with 8 $\mu g\ ml^{-1}$ polybrene. HaCaT keratinocytes and WTC11-hiPSCs were infected and then subjected to antibiotic selection (Hygromycin B, 50 $\mu g\ ml^{-1}$) for 5 days. For quadruple-sensor hiPSCs, an existing WTC11-AAV-CAG-GCaMP6f line was sequentially transduced with RFP-LifeAct (Puromycin 2 $\mu g\ ml^{-1}$) and FUCCIplex, then FACS-sorted to isolate double-positive clones (details in Supplemental Information).

*CRISPR/Cas9n knock-in at ROSA26.* To create a stable, two-color FUCCIplex reference line, the reporter cassette, flanked by 1 kb left/right homology arms for intron 1 of THUMPD3-AS1 (ROSA26), was PCR-amplified from pLV-EF1α-FUCCIplex. The donor fragment replaced EGFPd2 in pR26-Bsd_CAG-EGFPd2 via BamHI/MluI excision and NEBuilder HiFi assembly, generating pR26-Bsd_CAG-FUCCIplex. WTC11 hiPSCs were electroporated (1 pulse, 1200 V, 30 ms) with 1 µg donor plasmid plus paired Cas9 nickase vectors pSpCas9n(BB)_R26-L and -R, which introduce single-strand breaks flanking the insertion site. Following recovery, positive cells were clonally expanded. Correct integration was confirmed by 5′ and 3′ junction PCRs, absence of backbone PCR amplicons, and digital karyotyping (Illumina SNP array; no CNVs > 350 kb detected). Triply integrated clones (FUCCIplex + GCaMP6f + LifeAct) were produced by repeating the knock-in procedure in the double-sensor background.

*Cell culture.* HaCaT cells were maintained in DMEM/F12 without phenol red, supplemented with 10% FBS and 1% penicillin/streptomycin, splitting at 70% confluence to avoid contact inhibition. Human iPSCs (WTC11) were cultured on 1:100 growth-factor-reduced Matrigel in phenol-red-free Essential 8 Flex medium containing 1% P/S. At 70% confluence, they were dissociated for 5 min in 0.5 mM EDTA/DPBS at 37 °C, passaged 1:10 in the presence of RevitaCell, and fed every other day with fresh medium lacking RevitaCell.

*Directed cardiomyocyte differentiation.* Four days before induction, hiPSCs were seeded at $8\times10^4$ cells/well (Matrigel-coated 12-well plates) with RevitaCell. Day-0 (≈90 % confluence), the medium was switched to RPMI 1640 + GlutaMAX/B-27 minus insulin supplemented with 7.5 μM CHIR99021. On day-2, CHIR-supplemented medium was replaced with the same basal medium containing 2 μM Wnt-C59. Starting at day-4, a small molecule-free medium was exchanged, and from day-6, cells received cardio-culture medium (RPMI + B-27) every other day. Metabolic selection (day-12) employed glucose-free DMEM with 4 mM sodium DL-lactate for 4 days, after which hiPSC-CMs were maintained in cardio-culture medium. Beating cells were dissociated with 0.5 % trypsin-EDTA (10–15 min, 37 °C), counted, and replated at $5\times10^4$ cells/well on 0.2 % porcine-skin–coated ibidi μ-slides in cardio-culture medium plus 10 % FBS and 5 μM Y-27632. The medium was replaced the next day without FBS/Y-27632; calcium transient and sarcomere imaging were performed 5–7 days later.

*Cardioid first-heart-field organoids.* hiPSCs were seeded at $5 \times 10^4$ cells per well on Matrigel-coated 24-well plates in Essential 8 Flex + 5 μM Y-27632 (day-1). After 24 h, cells were switched to 1 ml anterior-primitive-streak (APS) induction medium for 36–40 h (day 1.5). Suspensions were aggregated in Nunclon Sphera U-bottom 96-well plates at $1.5 \times 10^4$ cells per well in CardMeso-ROCKi, centrifuged at 140 ×g for 4 min, and incubated for 24 h. The medium was changed daily with CardMeso (days 2.5–4.5) and then with CardMyo (days 5.5–6.5) until spontaneous beating emerged. From day 7.5, organoids were maintained in CardMaint medium refreshed every 48 hr. Media composition details are in the Supplemental Information file.

*Live-cell confocal imaging.* Confocal and wide-field experiments were performed on a Nikon inverted microscope equipped with a motorized 3-axis stage. Excitation was provided by a Lumencor Celesta light engine (405, 446, 477, 520, 546, 638, 740 nm; up to 800 mW) routed through multiband dichroics (Penta or CFP/YFP) and matching emission filters (e.g., 484/561 for mTurquoise2, 685/40 for miRFP-Geminin, 595/31 for RFP, 511/20 for EGFP). Fluorescence was captured on a Teledyne Photometrics Kinetix CMOS camera (6.5 μm pixels; 16-bit; 3 200 × 3 200 cropped to 2 700 × 2 700; no binning) through interchangeable air (20 × 0.75 NA) or silicone oil objectives (25× 1.05 NA, 40 × 1.25 NA, or 100 × 1.35 NA). Multichannel time-lapse or Z-stack sequences were acquired sequentially from 638 nm to 446 nm, with the shutter closed during stage or plane movements. Focus was maintained using Nikon Perfect Focus. The microscope was enclosed in an Okolab chamber maintained at 37 °C, 5% $CO_2$, and passive humidity; samples were imaged in phenol-red-free medium on #1.5H gelatin-coated coverslips or Ibidi/CuriBio plates. Image acquisition and hardware synchronization were managed in Nikon Elements AR v5.42.03, storing data as ND2 files.

*Live-cell light-sheet imaging.* For volumetric imaging, a Luxendo TrueLive3D light-sheet microscope was employed. It featured dual-scanned Gaussian illumination (sheet width ≈ 2 μm) and a 25 × 1.1 NA water-immersion objective coupled to twin Hamamatsu ORCA-Flash 4.0 V3 sCMOS cameras. Volumes were recorded using 445, 488, 561, and 642 nm lines; emitted light was filtered with band-pass filters: 418–462 nm for mTurquoise2 (or long-pass 466 nm), 580–627 nm for RFP, and 655–704 nm for miRFP. Samples were mounted in TruLive3D holders and maintained at 37 °C, 5 % $CO_2$ with active humidification in phenol-red-free medium. Volumes spanning 100 μm Z-range were captured at 0.5 μm steps, registered and fused in LuxBundle v5, and exported as H5 or TIFF stacks.

*FUCCIphase analysis.* The package accepts single-cell intensity tables exported from TrackMate (XML) or any CSV/Excel file that can be converted to a Pandas DataFrame, providing a unique cell identifier and paired nuclear intensities for mTurquoise2-Cdt1 and miRFP670-Geminin. Each intensity pair can be classified into G1, G1/S or S/G2/M using fixed thresholds as typically done with FUCCI datasets or aligned by dynamic time-warping to a reference trajectory recorded from 30 h HaCaT-FUCCIplex movies. By aligning a sequence of arbitrary length to the reference curve, FUCCIphase yields a continuous cell-cycle percentage per nucleus per frame. A 10-frame sequence led to a mean absolute error of ~20%. Including more frames improved accuracy. Optional utilities derive motility metrics from centroid coordinates or use TrackMate metrics to render phase-locked graphs in which any variable is plotted against the inferred percentage, rather than the frame number. The package is Made available on GitHub as open source (https://github.com/Synthetic-Physiology-Lab).*FUCCIsmart adaptive imaging.* The Nikon Elements JOBs macro (also on GitHub) utilizes basic intensity operations and can be easily adapted to different microscope control software. It surveys predefined fields every 6 hr by wide-field acquisition with 638 nm and 446 nm lasers through a 100× 1.35 NA silicone-oil objective; Perfect Focus maintains axial stability. Nuclear intensities corresponding to FUCCIphase> 37 % are flagged as S/G2/M, and those > 95 % as imminent mitosis. When an M-phase candidate is detected, the routine switches to spinning-disk mode, collects a 20 µm Z-stack at 2 µm steps in the LifeAct-RFP (546 nm) and EGFP-tubulin (477 nm) channels, and then resumes the survey loop. Shutters remain closed during stage or piezo movements to minimize phototoxicity. If no S/G2/M cells are present, the 6 hr cadence is retained; otherwise, the interval tightens to 1 hr until mitosis is captured, with the entire session limited to 24 hr. Across three independent runs, the workflow achieved > 65 % true-positive mitosis capture with < 8 % phototoxic events, as benchmarked against automated 1 h/6 h Z-stack controls. All acquisition parameters, thresholds and statistical analyses are detailed in Extended Figures 3–4.

*Reagents and working concentrations.* The SPY650-tubulin probe (SC503, Spirochrome) was dissolved in 50 µL of dimethyl sulfoxide (DMSO) to generate a 1,000× stock solution; hiPSC-CMs were then incubated for 1 hr at 37 °C with a 1× working solution immediately before live imaging. Nocodazole (M1404, 50 mg, Merck) was prepared at a concentration of 5 µg $\mu L^{-1}$ in DMSO. HiPSC-CM screens used a final concentration of 5 µM for 8 h, whereas hiPSC-derived ectoderm cultures received 100 nM and were imaged every 30 min for 18 h. For calcium-transient assays, HaCaT cells were loaded for 1 hr with 5 µM Fluo-4 AM (F14201, Thermo Fisher) and 0.025% (w/v) Pluronic F-127 (540025, Millipore) at 37 °C. Three PBS washes were followed by a 30-minute de-esterification in dye-free medium before imaging. Intracellular $Ca^{2+}$ release was evoked by a 100 µM ATP pulse during acquisition.

*Single-cell RNA sequencing.* To complement imaging readouts with molecular data, we performed single-cell RNA-seq on cardiac organoids. At day 6 of differentiation, organoids were dissociated with collagenase II and TrypLE. The 10x Genomics Chromium platform was used to capture 5,000–10,000 single cells per sample, followed by library preparation (Chromium Single Cell 3' v3 kit) and Illumina sequencing. The data were processed with Cell Ranger and analyzed in Seurat. Unsupervised clustering identified cardiomyocytes, proliferating cardiac progenitors, and other cell populations as classified by established scoring methods (Extended Fig. 8a). Raw and processed scRNA-seq data have been deposited in a public repository (https://www.ncbi.nlm.nih.gov/geo/query/acc.cgi?acc=GSE281528, token: wzilmgaglzqdzup), like the code (https://github.com/alessandro-bertero/FUCCIplex).

*Statistics.* Unless otherwise noted, values are reported as mean ± s.d. from at least three independent experiments. Sample sizes (number of cells, organoids, or imaging fields) for each analysis are given in the figure legends or SI. No statistical methods were used to predetermine sample size; experiments were replicated on different days or cell batches. Two-way ANOVA with post hoc Bonferroni test multi-group comparison or Kruskal-Wallis nonparametric with Dunn's test for multi-group comparison were used. Detailed statistical methods for each experiment can be found in the SI.
