## Supplementary information for "CALIPERS: Cell cycle-aware live imaging for phenotyping experiments and regeneration studies"

**Abstract:**

Cell cycle progression, migration, and proliferation shape development and regeneration, but simultaneous live-cell imaging remains challenging as conventional fluorescent cell cycle indicators (FUCCI) monopolize the green and red channels used by most structural and functional biosensors. To overcome this, we integrated a spectrally re-engineered FUCCI variant, open-source analysis software, and four-color human stem cell reporter lines into CALIPERS: a method for Cell-cycle-Aware Live-cell Imaging in Phenotyping and Regeneration Studies.

#### Supplementary Figures

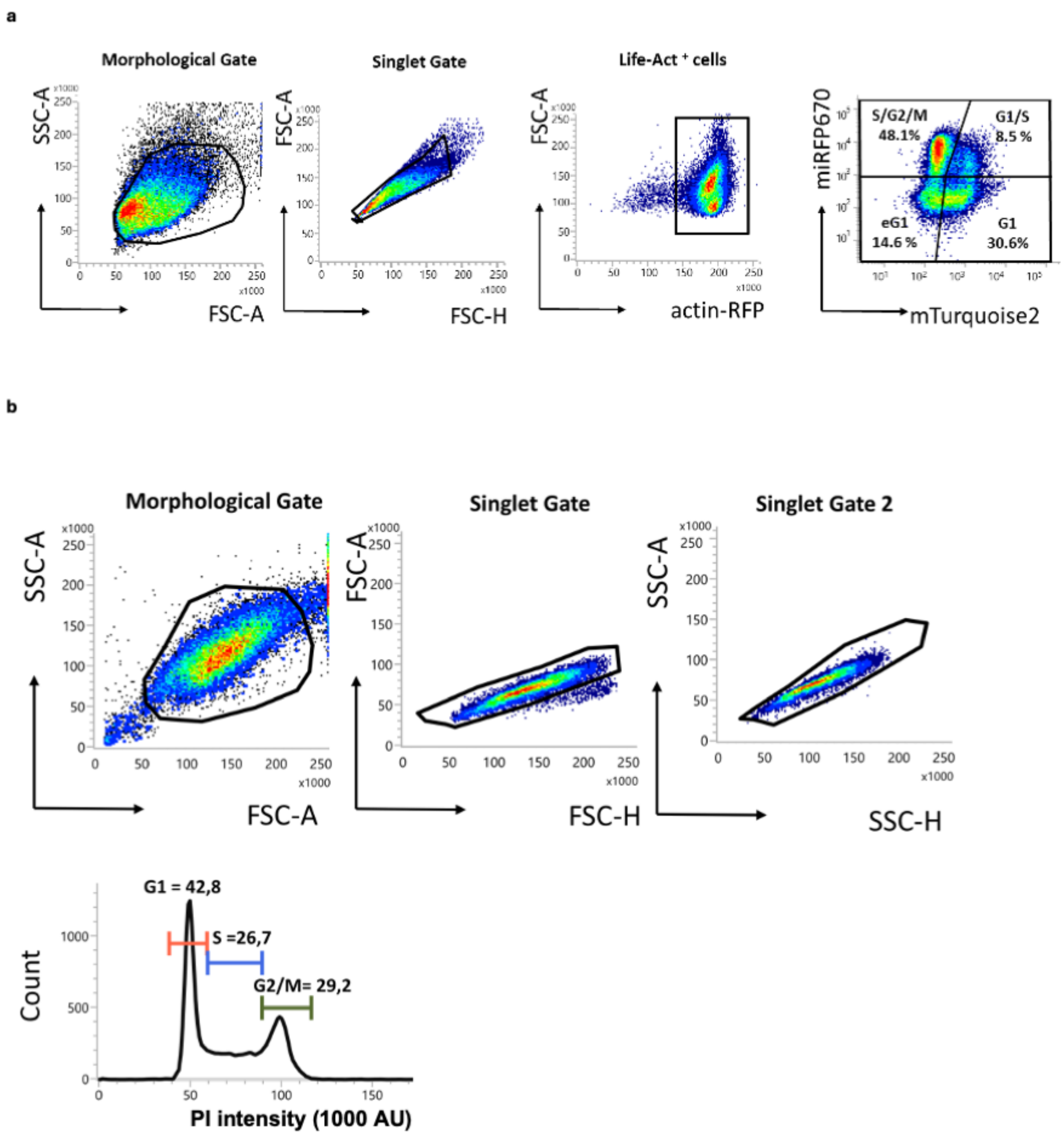


**Supplementary Figure 1. Cell cycle analysis of HaCaT-FUCCIplex cells**. **a)** Representative gating strategy in flow cytometry to identify the frequency of different cell subsets reflecting various phases of the cell cycle, based on RFP, miRFP670 and mTuqoise2 expression. The morphology gate and singlet cell gate exclude alive cells, debris, and doublets. **b)** Cell cycle analysis using propidium iodide (PI) staining and flow cytometry. The frequencies for the different cell cycle phases are indicated. Data is also shown in abbreviated form in the manuscript's main figure 1 and extended figure 1.

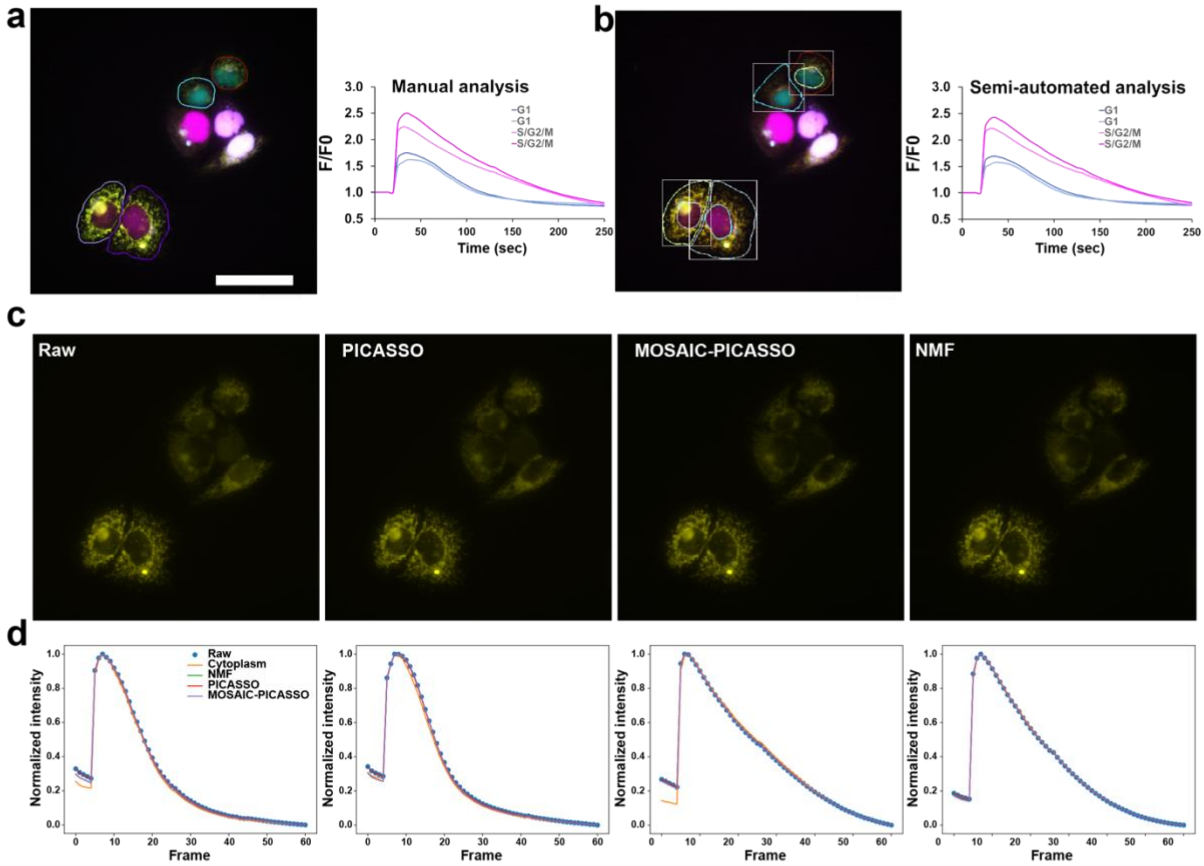


**Supplementary Figure 2: Dynamic multiplexing shown in HaCaT cells cycling calcium after an ATP dose. a-b)** Live imaging acquisition of calcium in HaCaT cells with FLUO-4 (yellow). The nuclear intensities determined the cell cycle phase (cyan - G1, magenta - S/G2/M). The intracellular calcium concentration was obtained from the average intensity inside cellular segmentation masks. The segmentation masks were either manually drawn (**a**) or obtained from Cellpose and manually curated using the frame with the highest calcium intensity (**b**). The normalized calcium concentration time courses obtained using the manual and semi-automated segmentation approaches are compared for selected cells. Data is also shown in abbreviated form in the manuscript's main figure 1. **c)** Representative frames comparing the effect of three bleed-through removal methods: PICASSO, MOSAIC-PICASSO, and NMF. The intensities were normalized by considering the maximum of the peak intensit. As the decay curves have almost identical slopes, the time dynamics (e.g., the time to reach 75% or 50% of the peak intensity) of the calcium signal are independent of the spectral overlap. **d)** Quantitative comparisons of cytoplasmic only calcium transients obtained from the raw images (blue dots), with a mask obtained by subtracting the nuclear mask from the cell mask to isolate the cytosplasm only (orange curve), and with the three bleedthrough removal methods mentioned above (green, red, and purple lines). All methods showed comparable results suggesting the G0/G1 bleed-through removal step is robust to method selection. Scale bar 25 μm.

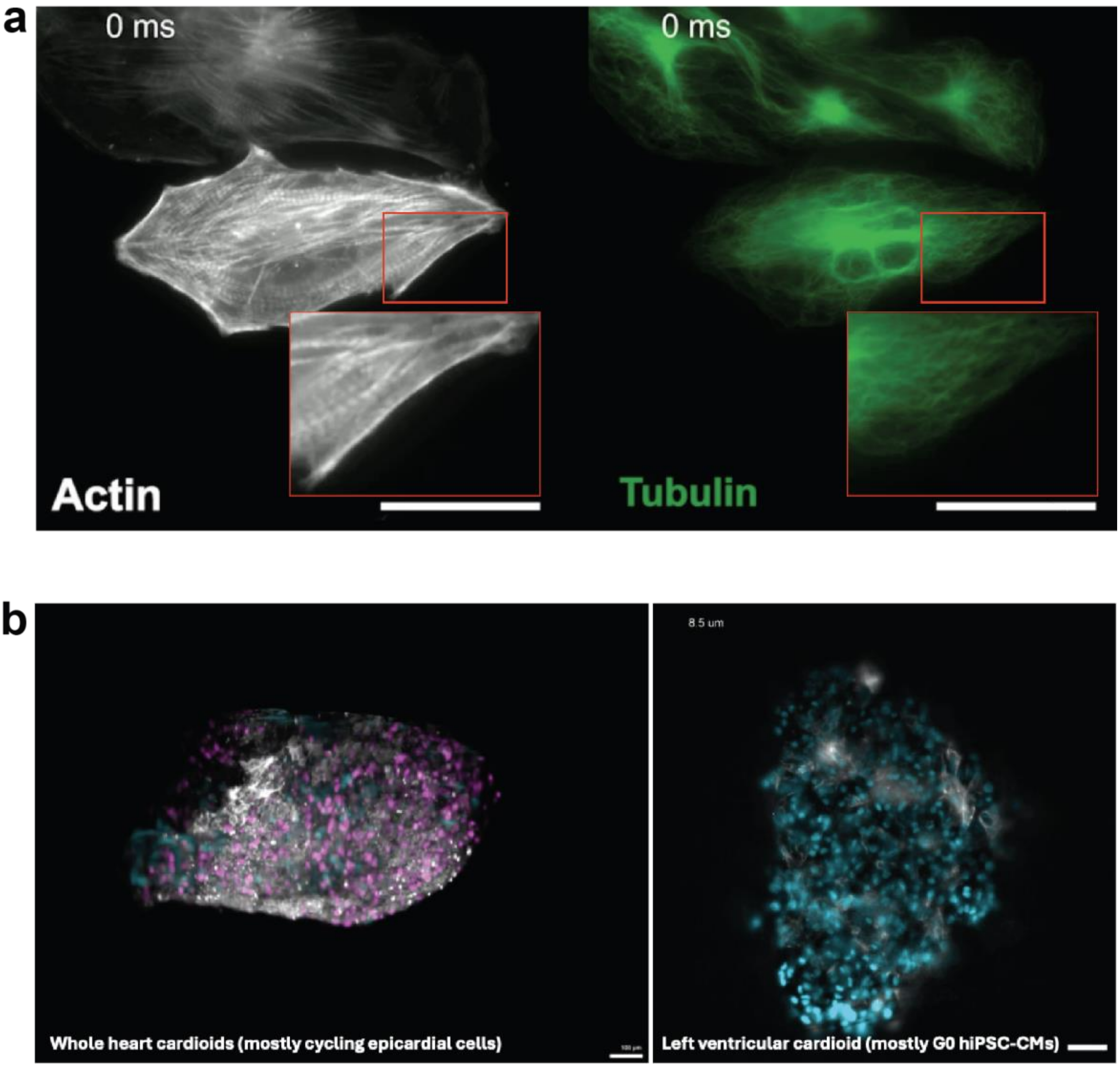


**Supplementary Figure 3: Comparison of the utilization of the clonal hiPSC-CMs obtained after lentiviral infection or genome editing. a**) An isolated frame from Supplementary Video 7 showing that FUCCIplex expression is silenced in LV-EF1a-FUCCIplex hiPSC-CMs, which enabled further multiplexing with a microtubule dye (Spyrochrome) to assess live sarcomere shortening and microtubule buckling simultaneously. **b**) An isolated frame from Supplementary Video 10 showing that FUCCIplex expression is retained in the CALIPERS hiPSCs (hR26-CAG-FUCCIplex), which were used to prepare whole heart (left) or left ventricular (right) cardiac organoids. In the former case, the outer layer exhibits numerous cycling epicardial cells. In contrast, the latter case is composed primarily of terminally differentiated hiPSC-CMs, characterized by constant nuclear CFP expression. Scale bars: 25 μm (a) 100 μm (b).

#### Extended Materials and Methods

##### *Molecular and cell biology*

**FUCCIplex design and production.** The FUCCIplex construct (pLV[Exp]-Hygro-EF1A-FUCCIplex) was obtained by swapping mKO2 and hmAzami Green from plasmid pBOB-EF1-FastFUCCI-Puro[1] (plasmid# #86849, Addgene) with mTurquoise2 (CFP) and miRFP670 (iRFP), respectively. An EF1a promoter drives the transgene expression. The new plasmid backbone is a third-generation mammalian gene expression lentiviral vector carrying a Hygromycin cassette controlled by an mPGK promoter. The generation of pLV[Exp]-Hygro-CMV-FUCCIplex, pLV[Exp]-Hygro-hTNNT2-FUCCIplex, and pLV[Exp]-Hygro-cTNT-FUCCIplex followed the same scheme, with the only exception that in these constructs, the expression of FUCCIplex is regulated by the listed promoters. Unless otherwise noted, Vector Builder conducted all sequencing and lentivirus packaging.

EF1α sequence[2,3] (length: 1179 bp):

GGCTCCGGTGCCCGTCAGTGGGCAGAGCGCACATCGCCCACAGTCCCCGAGAAGTTGGG
GGGAGGGGTCGGCAATTGAACCGGTGCCTAGAGAAGGTGGCGCGGGGTAAACTGGGAA
AGTGATGTCGTGTACTGGCTCCGCCTTTTTCCCGAGGGTGGGGGAGAACCGTATATAAGT
GCAGTAGTCGCCGTGAACGTTCTTTTTCGCAACGGGTTTGCCGCCAGAACACAGGTAAGT
GCCGTGTGTGGTTCCCGCGGGCCTGGCCTCTTTACGGGTTATGGCCCTTGCGTGCCTTGA
ATTACTTCCACCTGGCTGCAGTACGTGATTCTTGATCCCGAGCTTCGGGTTGGAAGTGGG
TGGGAGAGTTCGAGGCCTTGCGCTTAAGGAGCCCCTTCGCCTCGTGCTTGAGTTGAGGCC
TGGCCTGGGCGCTGGGGCCGCCGCGTGCGAATCTGGTGGCACCTTCGCGCCTGTCTCGCT
GCTTTCGATAAGTCTCTAGCCATTTAAAATTTTTGATGACCTGCTGCGACGCTTTTTTTCT
GGCAAGATAGTCTTGTAAATGCGGGCCAAGATCTGCACACTGGTATTTCGGTTTTTGGGG
CCGCGGGCGGCGACGGGGCCCGTGCGTCCCAGCGCACATGTTCGGCGAGGCGGGGCCTG
CGAGCGCGGCCACCGAGAATCGGACGGGGGTAGTCTCAAGCTGGCCGGCCTGCTCTGGT
GCCTGGTCTCGCGCCGCCGTGTATCGCCCCGCCCTGGGCGGCAAGGCTGGCCCGGTCGGC
ACCAGTTGCGTGAGCGGAAAGATGGCCGCTTCCCGGCCCTGCTGCAGGGAGCTCAAAAT
GGAGGACGCGGCGCTCGGGAGAGCGGGCGGGTGAGTCACCCACACAAAGGAAAAGGGC
CTTTCCGTCCTCAGCCGTCGCTTCATGTGACTCCACGGAGTACCGGGCGCCGTCCAGGCA
CCTCGATTAGTTCTCGAGCTTTTGGAGTACGTCGTCTTTAGGTTGGGGGGAGGGGTTTTAT
GCGATGGAGTTTCCCCACACTGAGTGGGTGGAGACTGAAGTTAGGCCAGCTTGGCACTT
GATGTAATTCTCCTTGGAATTTGCCCTTTTTGAGTTTGGATCTTGGTTCATTCTCAAGCCT
CAGACAGTGGTTCAAAGTTTTTTTCTTCCATTTCAGGTGTCGTGA

CMV sequence[4,5] (length: 589 bp):

TAGTTATTAATAGTAATCAATTACGGGGTCATTAGTTCATAGCCCATATATGGAGTTCCG
CGTTACATAACTTACGGTAAATGGCCCGCCTGGCTGACCGCCCAACGACCCCCGCCCATT
GACGTCAATAATGACGTATGTTCCCATAGTAACGCCAATAGGGACTTTCCATTGACGTCA
ATGGGTGGAGTATTTACGGTAAACTGCCCACTTGGCAGTACATCAAGTGTATCATATGCC
AAGTACGCCCCCTATTGACGTCAATGACGGTAAATGGCCCGCCTGGCATTATGCCCAGTA
CATGACCTTATGGGACTTTCCTACTTGGCAGTACATCTACGTATTAGTCATCGCTATTACC
ATGGTGATGCGGTTTTGGCAGTACATCAATGGGCGTGGATAGCGGTTTGACTCACGGGG
ATTTCCAAGTCTCCACCCCATTGACGTCAATGGGAGTTTGTTTTGGCACCAAAATCAACG
GGACTTTCCAAAATGTCGTAACAACTCCGCCCCATTGACGCAAATGGGCGGTAGGCGTGT
ACGGTGGGAGGTCTATATAAGCAGAGCTGGTTTAGTGAACCGTCAGATC

cTnT Sequence[6–8] (Length: 413 bp):

GGGATAAAAGCAGTCTGGGCTTTCACATGACAGCATCTGGGGCTGCGGCAGAGGGTCGG
GTCCGAAGCGCTGCCTTATCAGCGTCCCCAGCCCTGGGAGGTGACAGCTGGCTGGCTTGT
GTCAGCCCCTCGGGCACTCACGTATCTCCGTCCGACGGGTTTAAAATAGCAAAACTCTGA
GGCCACACAATAGCTTGGGCTTATATGGGCTCCTGTGGGGGAAGGGGGAGCACGGAGGG
GGCCGGGGCCGCTGCTGCCAAAATAGCAGCTCACAAGTGTTGCATTCCTCTCTGGGCGCC
GGGCACATTCCTGCTGGCTCTGCCCGCCCCGGGGTGGGCGCCGGGGGGACCTTAAAGCC
TCTGCCCCCCAAGGAGCCCTTCCCAGACAGCCGCCGGCACCCACCGCTCCGTGGGA

hTNNT2 promoter sequence[9] (length: 599 bp):

GTCATGGAGAAGACCCACCTTGCAGATGTCCTCACTGGGGCTGGCAGAGCCGGCAACCT
GCCTAAGGCTGCTCAGTCCATTAGGAGCCAGTAGCCTGGAAGATGTCTTTACCCCCAGCA
TCAGTTCAAGTGGAGCAGCACATAACTCTTGCCCTCTGCCTTCCAAGATTCTGGTGCTGA
GACTTATGGAGTGTCTTGGAGGTTGCCTTCTGCCCCCCAACCCTGCTCCCAGCTGGCCCTC
CCAGGCCTGGGTTGCTGGCCTCTGCTTTATCAGGATTCTCAAGAGGGACAGCTGGTTTAT
GTTGCATGACTGTTCCCTGCATATCTGCTCTGGTTTTAAATAGCTTATCTGAGCAGCTGGA
GGACCACATGGGCTTATATGGCGTGGGGTACATGTTCCTGTAGCCTTGTCCCTGGCACCT
GCCAAAATAGCAGCCAACACCCCCCACCCCCACCGCCATCCCCCTGCCCCACCCGTCCCC
TGTCGCACATTCCTCCCTCCGCAGGGCTGGCTCACCAGGCCCCAGCCCACATGCCTGCTT
AAAGCCCTCTCCATCCTCTGCCTCACCCAGTCCCCGCTGAGACTGAGCAGACGCCTCC

**HaCaT cell line culture and media.** HaCaT cells were kindly provided by Professor Hugo de Jonge (Department of Molecular Medicine, University of Pavia) and cultured in DMEM/F12 no phenol red (catalog# 21041-025, Gibco) with 10% Fetal Bovine Serum (FBS) (catalog # 10270-106, Gibco) and 1% Penicillin/Streptomycin (P/S) (catalog # A001, HiMedia). Cells were never allowed to reach full confluency and were split at 70% of confluence.

**Generation of the HaCaT FUCCIplex clonal cell line.** The generation of HaCaT FUCCIplex clonal cell line consisted of three different and sequential steps. First, HaCaT cells were infected with the FUCCIplex lentivirus and positively selected with antibiotics. A second lentiviral infection with a genetically encoded Lifeact-ACTB-RFP probe created a double reporter cell line that has the (F-)actin filaments tagged with a red fluorescent reporter in addition to the cell cycle indicator FUCCIplex. After antibiotic selection, cells were genome-edited to tag alpha-tubulin with an EGFP fluorophore at the N-terminus; this new 3-reporters cell line was then clonally expanded and selected. More in detail, HaCaT cells were plated at the density of $6 \times 10^4$ cells per well in a 12-well plate in complete DMEM/F12. The following day, the medium from the tissue culture plate was removed by aspiration and replaced with a fresh complete medium containing 8 µg/ml polybrene (VectorBuilder). Next, 10 µl of concentrated FUCCIplex lentivirus particles (>$10^8$ TU/ml, VectorBuilder) were resuspended in 500 µl of complete medium and added directly into the well. Cells were incubated overnight at 37 °C and 5% $CO_2$, and the next morning the transduction medium was aspirated and replaced with a fresh complete medium. After 4 days, positively transduced cells were selected with 20 µl/ml of Hygromycin B (50 mg/ml in PBS) (catalog # 10687010, Invitrogen). HaCaT FUCCIplex cells were then clonally seeded and expanded. Positive cells were evaluated by PCR and confocal imaging to assess the correct expression of the different reporters. For the Lifeact-ACTB-RFP lentivirus infection (rLV Ubi-LifeAct-TagRFP, $1 \times 10^7$ TU/ml, 10 µl), cells underwent the same procedure with the only difference that 0.62 µl/ml of Puromycin (1 µg/µl) was used to select positive cells. Finally, HaCaT cells were genome-edited using the ThermoFisher Scientific True Tag system. The donor sequence and primers with specific homology arms for GFP-tagging the N terminus of alpha-tubulin (TUBA1B, NM006082.3) were designed online with the assistance of the ThermoFisher Scientific TrueDesign Genome Editor tool. Briefly, the linear True Tag dsDNA donor oligo with the TUBA1A homology arms was amplified by PCR using Phusion Flash High-Fidelity PCR Master Mix, TrueTag Donor DNA Kit, GFP (catalog# A42992, Thermo). The TrueTag homology arm primer forward sequence is:

TCCTGTCGCCTTCGCCTCCTAATCCCTAGCCACTATGGGAGGTAAGCCCTTGCATTCG;

The reverse is:

CCTGAAAGCAGCCGGGAGCCGCACGGCTTACTCACACCGCTTCCACTACCTGAACC.

Next, the TrueGuide Synthetic gRNA (sgRNA) (ThermoFisher Scientific, sequence: GCACGGCTTACTCACCATAG) and True cut CAS9 Protein V2 (catalog# A36496, ThermoFisher Scientific) were combined to form the ribonucleoprotein (RNP) complex (CAS9+gRNA). True cut CAS9 Protein V2 (2 μg,12 pmol), sgRNA (400 ng, 12 pmol), True Tag donor (500 ng) were resuspended in 10 μl of Resuspension Buffer R (catalog# MPK1025, ThermoFisher Scientific) and then combined with 8 x $10^4$ cells resuspended in 8 μl of Resuspension Buffer R for a total final volume of 18 μl. 10 μl of this mix was used to electroporate the cells with the Neon transfection system (catalog# MPK5000, ThermoFisher Scientific). The conditions for electroporation were the following: 1200 V / 20 ms / 4 pulses. Electroporated cells were seeded in a 24-well plate filled with 0.5 ml of DMEM/F12 medium containing 10% FBS without antibiotics. The plate was incubated at 37 °C in a humidified $CO_2$ incubator. Positive cells were then clonally plated and expanded.

**hiPSCs maintenance, cardiomyocytes differentiation and selection.**

The WTC-11 (UCSFi001-A-1) hiPSC line and its derivative line carrying the GFP-based $Ca^{2+}$ indicator GCaMP6 (WTC11-AAV-CAG-GCaMP6) were kindly provided by Prof. Bruce Conklin (Gladstone. Institute of Cardiovascular Research). Cells were maintained in complete Essential 8 Flex Medium (catalog # A2858501, ThermoFisher Scientific) supplemented with Penicillin/Streptomycin (P/S) (catalog # A001, HiMedia) on 1:100 Growth Factor Reduced (GFR) Matrigel (7-10 mg/ml, depending on the lot), phenol red-free (catalog # 356231, Corning). When reached 70% confluency (usually in 3-4 days) cells were detached using 0.5 mM EDTA (Ethylenediaminetetraacetic acid, Life Technologies) in Dulbecco's PBS (DPBS) without $Ca^{2+}$ or $Mg^{2+}$ (catalog # C-40232, PromoCell) for 5 min at 37 °C and 5% $CO_2$ environment. Subsequently, cells were expanded in 6-well plates by passaging 1:10 in the presence of RevitaCell (catalog # A26445-01, LifeTechnologies). 24 h after plating, medium was replaced with complete Essential 8 Flex Medium (without RevitaCell) and changed every other day.

The hiPSCs were differentiated into cardiomyocytes (CMs) using a direct differentiation method adapted from the small molecule protocol[10]. More specifically, hiPSCs were plated at the density of 8 x $10^4$ /well in Matrigel pre-coated 12 well plates in the presence of RevitaCell three to four days before the beginning of the differentiation protocol (day -3 / -4). The day after, the medium was changed to complete Essential 8 Flex Medium (without RevitaCell). After 3-4 days, or when cells reached 90% of confluency (day 0), medium was replaced with pre-warmed cardio differentiation medium (RPMI 1640+ Glutamax Medium (1x), catalog # 72400, Gibco) supplemented with B-27 minus insulin (catalog # A1895601, Gibco) with the addition of 7.5 μM CHIR99021 (catalog # 72054, StemCell). Precisely after 48 h (day 2), medium was replaced with pre-warmed cardio differentiation medium with the addition of Wnt-C59 2 μM (catalog# 16644, Cayman Chemical Company). At day 4, when exactly 48 h elapsed, the medium was aspirated, and cardio differentiation medium was added. Cells were then left to differentiate for other 2 days. After 48 h (day 6), medium was changed and cells were fed with cardio culture medium (RPMI 1640+ Glutamax Medium (1x) plus B-27 Supplement (50x), serum-free (catalog # 17504044, Gibco)) and refreshed every 2 days. At day 12, cells were exposed to cardio Selection Medium (DMEM no Glucose (catalog #11966, Gibco) supplemented with 4 mM Sodium DL-lactate (catalog #71720, SIGMA) to metabolically select differentiated cardiomyocytes. The medium was refreshed every other day for 4 days, then cardiomyocytes were cultivated in the presence of cardio culture medium until experiments were performed following the specific time points of the assays. hiPSC-derived cardiomyocytes were detached with 0.5% of trypsin-EDTA solution (catalog# T3924) for 10-15 minutes at 37° C, counted and replated in a cardio culture medium supplemented with 10% FBS and Y-27632 dihydrochloride, 5 µM. Cells were plated into porcine skin-coated (0.2%, w/v) ibidi µ-slide 8 well-chambered coverslips (catalog # 80807, ibidi) at a density of 5 x $10^4$ cells/well. The day after, the medium was replaced with a fresh cardio culture medium without FBS and Y-27632.

**Generation of the hiPSC clonal cell line using lentiviral (LV) vectors (LV-EF1α-FUCCIplex).**

The strategy for generating hiPSC FUCCIplex clonal cell line followed the same framework adopted for creating the HaCaT FUCCIplex clonal cell line with some adaptation (see below).

To generate the double reporter (GCaMP6f and RFP-lifeAct) clonal cell line, the genetically encoded actin sensor RFP-LifeAct was introduced in the WTC11-AAV-CAG-GCaMP6 cell line. After antibiotic selection using 2µl/ml of Puromycin (1 µg/µl) positive clones were expanded. For the generation of the triple-reporter, 4-color hiPSC FUCCIplex clonal cell line, the FUCCIplex cell cycle indicator, and the genetically encoded actin sensor RFP-LifeAct were introduced in the WTC11-AAV-CAG-GCaMP6 cell line. After antibiotic selection using Hygromycin B (50 mg/ml in PBS) 50 µl/ml for FUCCIplex positive cells and 2µl/ml of Puromycin (1 µg/µl) for LifeAct positive cells, FUCCIplex hiPSc were sorted and the positive clones expanded.

**Generation of the CALIPERS hiPSC (hR26-CAG-FUCCIplex) clonal cell line**

To generate the single reporter, 2 color hiPSC HR26-CAG-FUCCIplex clonal cell line, the WTC-11 (UCSFi001-A-1) hiPSCs were genome-edited to specifically insert the FUCCIplex cassette into the ROSA26 locus via the THUMPD3-AS111 Site Usinga CRISPR/Cas9-Based gene-trap strategy. Unless otherwise indicated, genome editing was performed as previously described [12]. First, the FUCCIplex cDNA fragment flanked by the ROSA26 homology arms was amplified by PCR starting from the original FUCCIplex Plasmid (pLV-Hygro-EF1a-FUCCIplex) using primers fwd: CATCATTTTGGCAAAGAATTAATTCGGATCCgccaccatggtgagcaagg and rev: CGAGGCTGATCAGCGAGCTACGCGTttacagggccttccgccg. In parallel, the pR26-Bsd_CAG-EGFPd2 plasmid (kindly provided by Professor Alessandro Bertero, Department of Molecular Biotechnology and Health Sciences, University of Turin) was digested with BamHI and MluI (catalog

### R3136S and R3198S respectively, New England Biolabs) to remove the EGFPd2 fragment and ligated via NEBuilder HiFi DNA assembly (catalog# E5520S, New England Biolabs) with the FUCCIplex DNA fragment to generate the pR26-Bsd_CAG-FUCCIplex construct. Next, hiPSc were electroporated (1200V/30ms/1 pulse) in combination with other two plasmids, the pSpCas9n(BB)_HR26-L and pSpCas9n(BB)_HR26-R (kindly provided by Professor Alessandro Bertero) to induce a specific double-strand break in the intron between exons 1 and 2 of THUMPDS3-AS1 on chromosome 3 (ROSA26 locus). Afterward, cells were seeded in a 24-well plate without antibiotics and incubated at 37 °C in a humidified $CO_2$ incubator. Positive cells were then clonally plated and expanded. The strategy for generating the triple-reporter, 4-color HR26-CAG-FUCCIplex clonal cell line followed the same framework adopted for creating the single reporter, with the difference that the WTC-11 double reporter (GCaMP6f and RFP-LifeAct) was used: we called this line the CALIPERS hiPSC line. Gene-targeted hPSC clonal lines were screened by genomic PCR to verify site-specific targeting using the following primers:

| Locus | PCR type | | Primer | Sequence |
|---|---|---|---|---|
| ROSA | WT (locus) | 1 | FW | GAGAAGAGGCTGTGCTTCGG |
| | | 2 | REV | ACAGTACAAGCCAGTAATGGAG |
| ROSA | 5' INT | 3 | FW | GAGAAGAGGCTGTGCTTCGG |
| | | 4 | REV | AAGACCGCGAAGAGTTTGTCC |
| ROSA | 3' INT | 5 | FW | GAGAATAGCAGGCATGCTG |
| | | 6 | REV | ACAGTACAAGCCAGTAATGGAG |
| ROSA | 5' BB | 7 | FW | CGTTGTAAAACGACGGCCAG |
| | | 8 | REV | AAGACCGCGAAGAGTTTGTCC |
| ROSA | 3' BB | 9 | FW | GAGAATAGCAGGCATGCTG |
| | | 10 | REV | TGACCATGATTACGCCAAGC |

Wild Type (WT) ROSA 26 locus is detected by primers number 1 and 2. Site-specific cassette integration (INT) is detected by 5'INT and 3'INT PCRs (primers 3-4 and 5-6). Unintentional off-target random integrations were evaluated by two additional backbone (BB) PCRs: 5'BB and 3'BB (primers 7-8 and 9-10). PCRs were performed with Platinum PCR SuperMix High Fidelity (catalog # 12532016, Thermo), according to manufacturer instructions.

**Digital karyotyping:**

SNP-based human microarray was performed using genomic DNA by Life & Brain GmbH for Illumina Bead Array digital karyotyping. The data was analyzed with Genome Studio v2.0 (Illumina) to detect differences between the FUCCIplex-containing hiPSC lines and the parental lines. Copy number variations higher than $3.5x10^5$ bp and loss of heterozygosity higher than $1x10^6$ bp were considered significant.

**Lentiviral production and hiPSCs infection**

HEK293T cells were used for lentiviral production and maintained according to standard protocols. HEK293T cells were plated on 10 cm petri dishes at a density of 2.5 x $10^6$ cells/dish (day 0). Plates were coated with porcine skin gelatin solution, type A (0.2%, w/v) (catalog# G1890, Sigma). The day after (day 1), the lentiviral vector (FUCCIplex plasmid) and helper plasmids were transfected into the cells using Lipofectamine 2000 (catalog# 11668027, ThermoFisher Scientific). More specifically, 9 µg of FUCCIplex plasmid, 2.7 µg of envelope plasmid pMD2.G, 4.5 µg of pMDLg/pRRE plasmid and 1.8 µg packaging vector pRSV-Rev plasmid were resuspended in 1.5 ml of Opti-MEM (catalog# 31985070, ThermoFisher Scientific), mixed and added into 1,5 ml of Opti-MEM supplemented with 45 ul of Lipofectamine 2000. 12-16 hours post-transfection (day 2), media containing the DNA-lipofectamine 2000 complexes was replaced with 10 ml of DMEM (catalog# 41965062, Gibco). After 48 hours post-transfection (day 4) media with lentiviral particles was collected and centrifuged at 500g for 10 min, then filtered through 0.45 µm filter to remove cell debris. Lentiviral supernatant was concentrated with Lenti Concentrator (catalog# TR30025, Origene) following the manufacturer's instructions.

**Germ layers differentiation and Embryoid body formation**

For the direct differentiation of hiPSC into the 3 germ layers (endoderm, mesoderm, and ectoderm), the STEMdiff Trilineage Differentiation Kit (catalog# 05230, STEMCELL) was used according to the manufacturer's instructions. Briefly, for endoderm and ectoderm differentiation, 20 x $10^4$ cells were plated in an 8-well ibidi slide coated with 1:100 Growth Factor Reduced (GFR) Matrigel in E8 Flex medium supplemented with Y-27632 dihydrochloride, 5 µM. Conversely, 5 x $10^4$ cells were plated for mesoderm differentiation. The day after, the medium was replaced with the appropriate type of STEMdiff Trilineage media. For endoderm differentiation, cells were cultivated for 7 days. On the contrary, the mesoderm differentiation lasted for 5 days. Cells were imaged at the end of the differentiation.

To generate embryoid bodies, hiPSCs were detached, counted, and resuspended into Embryoid (EB) medium, consisting of DMEM/F12+GlutaMAX (catalog# 10565-018, Thermo), KnockOut SR (catalog# 10828-028, Thermo), Non-Essential Amino Acids (catalog# 11140-076, Thermo) supplemented with 2-mercaptoethanol (catalog# 31350010, Thermo) for a 0.1 mM final concentration. Cells were seeded at a final concentration of 1 x $10^4$ cells per well (Nunclon sphera 96-Well, U-Shaped-

Bottom Microplate, catalog#174929, Thermo) in the presence of Y-27632 dihydrochloride, 5 µM (catalog # 004CA16644, Cayman Chemicals). Afterward, cells were centrifuged at 140 xg for 4 minutes, incubated at 37 °C, and 5% $CO_2$ for 24h. The day after, the entire medium was changed and replaced with EB medium without Y-27632 dihydrochloride. EBs medium was changed every other, for 14 days, without disturbing the embryoid bodies.

**Generation of 3D self-assembling human heart organoids (First Heart Field)**

To generate cardiac organoids (cardioids) from CALIPERS hiPSCs, we used the protocol previously described[20,21] with minor changes. Briefly, for the anterior primitive streak (APS) induction (day -1), CALIPERS hiPSC cells were seeded at 5 x $10^4$ cells per well (Matrigel-coated 24 well plate) in the presence of Essential 8 Flex + Y-27632 dihydrochloride, 5 µM (catalog # 004CA16644, Cayman Chemicals). After 24h, the supernatant was removed and replaced with 1 ml APS- induction medium per well (see table below) and cells were incubated at 37 °C and 5% $CO_2$ for 36-40h. On day 1.5, cells were detached, counted, and resuspended in CardMeso-Rocki (see table below) to a final concentration of 1,5 x $10^4$ cells per well (Nunclon sphera 96-Well, U-Shaped-Bottom Microplate, catalog#174929, Thermo). Cells were centrifuged at 140 xg for 4 minutes and incubated at 37 °C and 5% $CO_2$ for 24h. From day 2.5 to 4.5, the medium was changed every day and wells were filled with CardMeso-induction medium without ROCKi. From day 5.5 to 6.5 the medium was exchanged for CardMyo-induction medium (see table below). Organoids started beating at this stage. From day 7.5, the supernatant was exchanged for CardMaint medium (see table below) and replaced every 48h.

| **Reagent name** | **Supplier and catalog number #** |
|---|---|
| Bovine serum albumin (BSA) | Europa Biosciences, #EQBAH70-1000, Lot #190618-0170 |
| Optiferrin | Invitria, #777TRF029 |
| Human FGF2-G3 | Qkine, #Qk053 |
| Activin A1 | Qkine, #Qk001 |
| BMP4 | Qkine, #Qk038 |
| Insulin | Thermo Scientific, #A11382II |
| Y-27632 dihydrochloride (ROCK inhibitor, ROCKi) | Cayman Chemicals, #004CA16644 |
| LY 294002 hydrochloride | Selleckchem, #1105 |
| CHIR99021 | Cayman Chemicals, #004CA13122 |
| C59 | Cayman Chemicals, #16644 |
| Retinoic acid | Sigma Aldrich, #R2625 |

| | |
|---|---|
| F-12 | Thermo Scientific, #31765068 |
| IMDM | Thermo Scientific, #21980032 |
| Chemically defined lipid concentrate | Thermo Scientific, #11905031 |
| Mono-Thio-Glycerol (MTG) | Sigma Aldrich, #M6145 |

| Medium | Reagent | Final Concentration |
|---|---|---|
| **CDM** | BSA<br>IMDM<br>F-12<br>Conc. lipids<br>MTG<br>Optiferrin | 0.3 %<br>~50%<br>~50%<br>1%<br>0.004%<br>15µg/ml |
| **APS – Induction (CDM-FLyABCh)** | **CDM**<br>FGF2-G3<br>LY 294002<br>Activin A1<br>BMP4<br>CHIR99021 | <br>6ng/ml<br>5µM<br>5ng/ml<br>10ng/ml<br>5µM |
| **CardMeso – Induction (CDM-BFIC59Ra)** | **CDM**<br>BMP4<br>FGF2-G3<br>Insulin<br>C59<br>RA<br>**Y-27632**<br>**Day 1.5 only** | <br>10ng/ml<br>1.6ng/ml<br>10µg/ml<br>2µM<br>50nM<br>**5µM** |
| **CardMyo – Induction (CDM-BFI)** | **CDM**<br>BMP4<br>FGF2-G3<br>Insulin | <br>10ng/ml<br>1.6ng/ml<br>10µg/ml |
| **CardMaintenance (CDM-I)** | **CDM**<br>Insulin | <br>10µg/ml |

**Generation of self-assembling human heart organoids (hHOs)**

To generate human heart organoids (hHOs) we used the protocol previously described[22] with minor changes. Specifically, to generate embryoid bodies (day-2), human pluripotent stem cells (hPSCs) were maintained at a sub-confluent density (60-80% confluence) and washed with Dulbecco's Phosphate-Buffered Saline (DPBS). Following this, 1 mL of PBS/0.5mM EDTA was added to each well for 3-6 minutes. The dissociation was halted by adding 1 mL of Essential 8 Flex medium supplemented with

RevitaCell (100X). Cells were counted and resuspended at a concentration of 100,000 cells/mL. Subsequently, 100 µL (10,000 cells) of the cell suspension was added to each well of a round-bottom ultra-low attachment 96-well plate. The plate was centrifuged at 100 × g for 3 minutes and incubated for 24 hours at 37 °C in a 5% CO2 atmosphere. On Day -1, 50 µL of the existing medium was carefully aspirated from each well of the culture plate, followed by the addition of 200 µL of fresh Essential 8 Flex medium pre-warmed to 37 °C, resulting in a total volume of 250 µL per well. The cells were then incubated for 24 hours at 37 °C in a 5% CO2 atmosphere. On Day 0, to initiate differentiation towards the mesoderm lineage, 166 µL of medium was removed from each well. Subsequently, 166 µL of RPMI 1640 medium supplemented with insulin-free B-27 was added, along with the following reagents: 6 µM CHIR99021, 1.5 ng/mL Activin A, and 1.875 ng/mL bone morphogenetic protein 4 (BMP4). After 24 hours of incubation at 37 °C and 5% CO2, the medium was replaced again by removing 166 µL and adding fresh RPMI 1640 with insulin-free B-27.On Day 2, cardiac mesoderm specification was induced by removing 166 µL of the medium and adding 166 µL of RPMI 1640 supplemented with insulin-free B-27 and 3 µM Wnt-C59, resulting in a final concentration of 2 µM Wnt-C59. The cells were incubated for 48 hours under the same conditions. On Day 4, the medium was refreshed again by removing 166 µL and adding 166 µL of RPMI 1640 with insulin-free B-27, followed by another 48-hour incubation.

On Day 6, the medium was replaced with RPMI 1640 supplemented with B-27 and insulin, and the cells were incubated for 24 hours. On Day 7, to promote pro-epicardial differentiation, an initial volume of 166 µL of medium was aspirated from each well. Subsequently, 166 µL of fresh RPMI 1640 medium supplemented with B-27 and insulin, along with 3 µM CHIR99021, was added, achieving a final concentration of 2 µM CHIR99021 in each well. The cells were then incubated for one hour at 37°. Afterwards, 166 µL of medium from each well was removed and 166 µL of fresh RPMI 1640 containing B-27 supplement (+ insulin) was added. See also Fig. S3 and Supplementary video 10 for a comparison of the two types of cardiac organoid platforms.

**Flow Cytometry and Cell cycle analysis**

For flow cytometry analysis, HaCaT cells were detached with Trypsin/EDTA 0.25% (catalog# 25200056, Thermo), resuspended in culture medium supplemented with 10% of FBS and pelleted at 300 g for 5 minutes. Subsequently, cells were washed twice with PBS (without $CaCl_2/MgCl_2$), pelleted, and resuspended at the density of 5 x $10^5$ cells per ml. Before analysis, cell clusters were disaggregated using the FACS flow tube caps (catalog# 352235) containing a 35µm nylon mesh screen. For cell cycle analysis, 5 x $10^5$ cells were detached with Trypsin/EDTA 0.25%, pelleted at 300 g for 5 minutes, washed twice with PBS (without $CaCl_2/MgCl_2$), resuspended and fixed with cold ETOH 70%. After 30 minutes at 4 degrees, cells were pelleted at 850 g for 5 minutes, washed twice with PBS (without

$CaCl_2/MgCl_2$), and stained with PBS-propidium Iodide (PI) (catalog# P 4170, Sigma) and RNase A (catalog# EN0531, Thermo ) at a final concentration of 0.05 and 0.1 mg/mL, respectively. At least 2 x $10^4$ unicellular events were analyzed for each experimental condition. All the flow cytometry experiments were conducted with FACSLyric (BD, Franklin Lakes, NJ, USA) equipped with three lasers (405, 488, and 640 nm), and analyzed by FlowJo software (BD,Biosciences). For the cell cycle, the analysis was performed with "Cell Cycle" plugin of Flow Jo. See also Fig. S1.

**Sorting**

For Flow Cytometry Activated Cell sorting (FACS), cells were detached with Trypsin/EDTA 0.25%, resuspended in culture medium supplemented with 10% of FBS and pelleted at 300 xg for 5 minutes. Subsequently, cells were washed twice with PBS (without $CaCl_2/MgCl_2$), pelleted, and resuspended at the density of 2 x $10^6$ cells per ml in PBS, filtered with FACS flow tube caps (catalog# 352235) containing a 35µm nylon mesh screen ml and acquired. FACS was performed on a Becton Dickinson FACS Aria III flow cytometer equipped with four lasers (405, 488, 571 and 640 nm). Phenotypic analysis and gating strategy were performed with BD FACS Diva software. Cells were separated with a 100 µm nozzle, at room temperature and with a low flow rate. Single cell sorting was performed in 96-well plate; index sorting analysis was performed at the end of the sorting experiment. Cells after the sorting, were immediately used for experiments. See also Fig. S1.

***Imaging and image analysis***

**Widefield and confocal imaging**

Imaging was performed using a fully motorized Nikon Ti-2 inverted widefield microscope equipped with the Perfect Focus System and a Crest V3 X-Light spinning disk confocal module. The microscope featured a Nikon linear-encoder motorized stage and a Mad City Labs 100 µm-range Z piezo insert (catalog #NI-2-C312, Mad City Labs). Fluorescence excitation was provided by a Lumencor Celesta Light Engine (405, 477, 520, 546, 577, 638, and 740 nm; catalog #TSX5030FV), delivering up to 800 mW of laser power at the fiber tip via 7 solid-state laser lines. In this study, we used directly modulated solid-state lasers at 638, 546, 477, and 446 nm. Excitation light passed through a dichroic mirror wheel containing either a hard-coated Full Multi-Band Penta CELESTA -DA/FI/TR/Cy5/Cy7-A dichroic (catalog #MXR00543, Nikon) or a Full Multiband Dual CELESTA -CFP/YFP-A (catalog #MXR00544, Nikon). Depending on the laser line, light was subsequently filtered through either a Multiband Full Penta FF01-391/477/549/639/741 (catalog #FL-416877, Semrock) or a Full Multiband Dual FF01-449/520 (catalog #FL-411981, Semrock). Emitted fluorescence was collected using a Teledyne Photometrics Kinetix CMOS camera and a selection of Nikon objective lenses matched to the experiment, including CFI Plan Apo 20× (air-immersion, N.A. 0.75, WD 1 mm, catalog #MRD00205), CFI Plan Apo Lambda S 25×C Sil (oil-immersion, N.A. 1.05, WD 0.55 mm, catalog #MRD73250), CFI Plan Apo Lambda S 40×C Sil (oil-immersion, N.A. 1.25, WD 0.3 mm, catalog #MRD73400), and CFI SR HP Plan Apo Lambda S 100×C Sil (oil-immersion, N.A. 1.35, WD 0.31–0.28 mm, catalog #MRD73950). Silicon immersion oil (Nikon, 30 cc, catalog #MXA22179) was used as required.

Emission light was collected through one of the designated filters in the emission wheel: FF01-484/561 (catalog #FL-412124, Semrock) for mTurquoise2, FF01-685/40-25 nm (catalog #FL-011482, Semrock) for miRFP, FF01-595/31 (catalog #FL-004391, Semrock) for RFP, and FF01-511/20-25 (catalog #FL-004306, Semrock) for EGFP. Channels were acquired sequentially, following a far-red (638 nm) to cyan (446 nm) laser line order. Images were captured at 3200 × 3200 pixels and cropped to 2700 × 2700 pixels (6.5 µm pixel size) using a 16-bit digitizer without binning. Acquisition was managed via Nikon Elements AR NIS software (version 5.42.03), and meta-/data were saved in ND2 format.

All experiments were conducted under live imaging conditions using a phenol red-free imaging medium. Samples were prepared on #1.5H glass coverslips coated with 0.2% porcine gelatine (catalog #G2625-500G, Sigma-Aldrich) in round dishes, 8-well plates, 96-well formats (catalog #81158, #80807, #89626, Ibidi), or 96-well nanosurface plates (catalog #ANFS-0096, CuriBio). The microscope was housed in an Okolab enclosure to maintain physiological conditions at 37 °C with 5% $CO_2$ and passive humidification. Confocal and widefield imaging were both employed as needed. Experiments typically included sequential multichannel acquisitions, multiple fields of view (FOV), and time-lapse imaging. The shutter remained closed during stage movements or idle periods, and the Perfect Focus System was used to maintain focus. For Z-stack imaging, stacks were acquired using the Mad City Lab (MCL) Piezo Z-device, either by capturing all channels across the full Z-stack or each focal plane across all channels.

**Light-sheet imaging**

Light-sheet imaging was performed using the Luxendo TruLive3D microscope, which employs dual-scanned Gaussian beam illumination and four discrete laser lines (446, 488, 561, and 642 nm) with a selected light-sheet width of 2 µm. Fluorescence detection was achieved using a 25× NA 1.1 water-immersion objective and two Hamamatsu sCMOS Orca Flash 4.0 V3 cameras. Static volumetric imaging was conducted using all four laser lines to capture distinct self-assembly stages of the cardioids. Emitted light was filtered using a 418–462 nm band-pass filter for mTurquoise2, a 655–704 nm band-pass filter for the Geminin signal, and a 580–627 nm band-pass filter for RFP. Channels were acquired sequentially, following a far-red (638 nm) to cyan (488 nm or 446 nm) order. Image acquisition was controlled via LuxBundle software (version 5), with subsequent registration and fusion also performed within the platform. Data were saved in both H5 and TIF formats.
All experiments were performed under live imaging conditions using phenol red-free imaging medium. Samples were mounted using the Luxendo TruLive3D sample holder. The microscope's enclosed sample chamber maintained physiological conditions at 37 °C with 5% $CO_2$ and active humidification. Volumetric datasets were acquired with a 0.5 µm step size across a 100 µm z-range, enabling high-resolution, multi-channel 3D visualization of cardiac morphogenesis.

**Structural and functional phenotyping in HaCaT cells and hiPSC-CMs**

To monitor cytosolic calcium transients, HaCaT cells were seeded at a density of $25 \times 10^3$ cells/well in 0.2% (w/v) porcine skin-coated µ-slide 8-well chambered coverslips (catalog #80807, Ibidi). The following day, cells were incubated in DMEM-F12 containing 5 µM Fluo-4 AM (catalog #F14201, ThermoFisher Scientific) and 0.025% (w/v) Pluronic F-127 (catalog #540025, Millipore) for 1 h at 37 °C in the dark. After three PBS washes, the cells were incubated in DMEM-F12 for 30 minutes to ensure complete de-esterification of intracellular AM esters. Optical recordings were performed in widefield mode using a Crest V3 X-Light spinning disk confocal microscope (Nikon) equipped with a Celesta Light Engine (catalog #TSX5030FV, Lumencore), a Photometrics Kinetix sCMOS camera, and a Nikon CFI SR HP Plan Apo Lambda S 100XC Sil objective (oil-immersion, N.A. 1.35, WD 0.31–0.28 mm; catalog #MRD73950, Nikon). For calcium transient induction, ATP was applied at 100 µM. Cells were sequentially illuminated at 638, 546, 477, and 446 nm with 30 ms exposure every 5 seconds for a total duration of 5 minutes. Calcium imaging was acquired with the 477 nm laser line. Raw data were analyzed in ImageJ. At least three wells per sample were imaged across five independent experiments. For hiPSC-CMs we followed a similar analysis: after 5-7 days after the end of differentiation (see above), calcium transients were acquired with a 477nm laser line with an exposure time of 30 ms. For $Ca_{2+}$ imaging, traces were analyzed using ImageJ's Time Series Analyzer tool, and data were expressed as ΔF/F0 or ΔF/(Fmax-F0) as needed. For sarcomere profile analysis, cardiomyocytes were imaged using a 546 nm laser line to acquire information on the contractile structure organization, and the images were analyzed with ImageJ using the plot profile tool.

**Automated elimination of the bleed-through artifacts from CFP-expressing nuclei**

HaCaT cells were illuminated in sequence at 638, 546, 477, and 446 nm with an exposure time of 30 ms every 5 seconds for 5 minutes (100x magnification). Actin, calcium, and the nuclear FUCCIplex channels (cyan and magenta) were recorded. Since cells and nuclei did not move significantly during the acquisition. The first frame was used to segment the nuclear channels. Each mask was assigned a cell cycle phase (G1, G1/S or S/G2/M) based on the cyan and magenta intensity. The frame with the highest mean calcium intensity was used to prepare segmentation masks to measure the calcium intensity. We used Cellpose v2.0[13] to segment the calcium channel. The nuclei were segmented as used in the other analyses using StarDist[14] on a merged nuclear channel. All segmentation masks were manually curated as necessary. Segmentation masks touching the frame border were excluded. Then, nuclei were mapped on cells and cell masks with multiple nuclei were excluded. Nuclei with a strong cyan signal were visible in the calcium channel due to spectral overlap. To address this issue, we explored the following unmixing techniques: non-negative matrix factorization (NMF)[15], PICASSO and MOSAIC-PICASSO[16–18]. In addition, we considered only the extranuclear calcium signal by subtracting the nuclear calcium signal. All techniques were implemented in Python. We leveraged the NMF implementation in scikit-learn[19], the Napari-PICASSO implementation (https://github.com/nygctech/PICASSO), and the MOSAIC-PICASSO package accompanying the original study (https://github.com/xing-lab-pitt/mosaic-picasso). To determine if the spectral overlap affects the dynamics of the calcium signals, we considered the normalized curves, as the unmixing techniques do not yield consistent absolute intensity values. We found that the dynamics of the calcium signal decay, e.g., the time to reach 50% of the maximum intensity, were not impacted by the spectral overlap (see an example in Fig. S3). The example shown in Fig. 1 of the main text was unmixed using NMF.

**FUCCIphase analysis.**

We developed a Python package to facilitate the analysis of the cell cycle based on the single-cell FUCCI information. Supported input formats are TrackMate[23] XML files or Pandas DataFrame objects, which can be initialized from CSV or Excel files. Information about the FUCCI channel intensities and a unique cell identifier needs to be present. The channel intensity pair for each cell was associated with a cell cycle phase, i.e. G1, G1/S or S/G2/M in the case of the FUCCI sensor considered here. The software layout supports, in general, also other FUCCI sensors, but they have not yet been implemented.

FUCCIphase permits to reconstruct the cell cycle percentage of individual cells (see Fig. 1, Ext. Fig. 2). We collected a reference intensity curve for the entire cell cycle of HaCaT cells from live cell recordings over 30 hours (see Ext. Fig. 2b). The cell cycle percentage was obtained by dividing the time of the cell cycle by the total time of the cell cycle. First, we attempted to reconstruct the cell cycle percentage by a look up of the cyan and magenta intensity pair on the reference curve pair. This approach yielded limited accuracy. Instead, we estimated the cell cycle percentage by using a sequence of channel intensity pairs. The sequence is used to find a matching subsequence of the reference curve (see Ext. Fig. 2c). As a consistency test, we matched the full cycles used to distill the reference curve against the reference curve. Here, the cell cycle percentage as ground truth is known and we could observe a very good agreement between ground truth and reconstructed cell cycle percentage (Ext. Fig. 2d). We then sampled subsequences of varying length and tested the accuracy of the reconstruction (Ext. Fig. 2e). A subsequence length of 10 frames permits on average a percentage estimate error of less than 20%.

FUCCIphase features utilities to compute motility parameters from the input data if information about the nucleus centroids is present (see Fig. 1g). Furthermore, features computed by TrackMate can be considered. FUCCIphase offers visualization routines to plot phase-locked parameters, i.e., parameters are plotted over the estimated cell cycle percentage instead of the frame number or time (see Ext. Fig. 2f-i). In addition, it interfaces the LineageTree library (https://github.com/GuignardLab/LineageTree) to plot lineage trees (Fig. 1g, Ext. Fig. 2a). We interfaced Napari[24] and provide routines to overlay the fluorescence microscopy images with estimated cell cycle percentages and labels coloured according to the cell cycle.

**FUCCIsmart Routine**

The FUCCIsmart routine (available at github.com/Synthetic-Physiology-Lab) automates the switch from 2D to 3D imaging upon mitotic entry, marked by the cytoskeletal transition from a flat microtubule network to a 3D mitotic spindle. Developed in Nikon Elements AR NIS software using the JOBs module, FUCCIsmart was tested on HaCaT FUCCIplex cells. Cell cycle classification was performed using FUCCIphase, which computes % cell cycle completion based on FUCCI fluorescence intensity. Cells exceeding 37% were identified as S/G2/M, and those above 95% were entering mitosis.

The screening was performed via widefield imaging using 638 nm and 446 nm lasers and a Nikon CFI SR HP Plan Apo Lambda S 100XC Sil objective (oil-immersion, N.A. 1.35, WD 0.31–0.28 mm; catalog #MRD73950, Nikon). When M-phase entry was detected, the system triggered a hardware switch to spinning disk confocal mode for live 3D imaging. Z-stacks were acquired using an MCL Piezo Z-device with 2 µm steps across a 20 µm range, capturing Lifeact-ACTB-RFP (546 nm) and EGFP-Tubulin (477 nm) channels using the same objective. The routine operated over 24 h under live imaging conditions, merging sequential multichannel acquisition, Z-stacks, and nested time-lapse imaging. If no S/G2 cells were present, screening was performed every 6 h. Upon S/G2 detection, FUCCIsmart increased screening frequency to once per hour to avoid missing the 95% mitotic threshold. Hourly imaging is triggered when a cell enters S/G2/M (FUCCI phase> 37%) to avoid missing the 95% mitosis threshold. Two acquisition modes were implemented: Single FUCCIsmart, which captured a single Z-stack at mitosis onset, and Multi FUCCIsmart, which acquired Z-stacks every 15 min for 1 h to capture mitotic progression.

HaCaT cells were seeded at $15 \times 10^3$ per well on #1.5H glass coverslips coated with 20 µg/mL fibronectin (catalog #341635-1MG, Sigma-Aldrich; #81158, Ibidi). Cells were incubated overnight (15 h) to allow spreading and re-entry into the cell cycle. Before imaging, medium was replaced with DMEM/F12 without phenol red (catalog #21041-025, Gibco), supplemented with 10% FBS (catalog #10270-106, Gibco) and 1% Pen/Strep (catalog #A001, HiMedia). Imaging was conducted at 37 °C with 5% $CO_2$ and passive humidification. Control conditions included 24 h experiments with automated Z-stacks every 1 h or 6 h. All data were saved as .nd2 files, with each Cell ID containing FOV coordinates, time-loop index, cell cycle class, nested loop index, and the 'Mcell' flag if Z-stacks were acquired. These logs enabled full reconstruction of cell fate and phototoxicity analysis. FUCCIsmart outputs were classified as True/False Positive or True/False Negative based on visual validation (Extended Fig. 3). Performance metrics were benchmarked across 3 independent experiments and compared to controls (Extended Fig. 4), using FUCCIphase-estimated cell cycle durations (G1: 9 ± 2.3 h; S/G2/M: 10 ± 1.15 h). Phototoxic events were flagged when cells died or showed abnormal phase elongation and were marked with a black box in the classification table. The percentage of mitoses acquired and phototoxicity events in the Single and Multi FUCCIsmart routines were compared against automated Z-stack acquisitions performed every 1 h and 6 h (positive controls). Mitosis acquisition rate was calculated over the initial number of cells, while phototoxicity rate was calculated relative to the expected number of progeny at 24 h (expected twice as many cells as the original number).

Each condition was tested in $n = 3$ independent experiments. Fields of view (FOVs) with imaging artifacts or cell loss were excluded from analysis. All values are reported as mean ± standard deviation (SD). Statistical analysis was performed using GraphPad Prism (v10.3.01) and SigmaPlot (v14.0). A two-way ANOVA was used to assess variation between groups and readouts (mitosis acquisition and phototoxicity). Multiple comparisons were corrected using Bonferroni's method. Significance was defined as $p < 0.05$. Significant comparisons included (Figure 1i):

Mitosis Acquired (%)

- FUCCIsmart Single vs. Automation 6 h: $p < 0.0001$
- FUCCIsmart Multi vs. Automation 1 h: $p = 0.0042$
- FUCCIsmart Multi vs. Automation 6 h: $p = 0.014$
- Automation 6 h vs. Automation 1 h: $p < 0.0001$

Phototoxic Events (%)

- FUCCIsmart Single vs. Automation 1 h: $p < 0.001$
- FUCCIsmart Multi vs. Automation 1 h: $p < 0.0001$
- Automation 6 h vs. Automation 1 h: $p < 0.0001$

Full datasets and statistical outputs are provided in the Data availability section.

**CALIPERS in cardioids**

Light-sheet datasets were processed using ARIVIS software with a dedicated pre-processing pipeline. Three-channel volumes were background-corrected, and a median filter was applied to the nuclear channels. Lucy-Richardson deconvolution (10 iterations) was performed on the actin channel, followed by denoising. Nuclear segmentation in the mTurquoise2 and miRFP670 channels was performed using a deep-learning model in ARIVIS Cloud. Segmentation was performed on cropped sub-volumes of 25 μm depth for both nuclear channels at each time point. Detected objects were quantified to estimate the percentage of cells expressing mTurquoise2 and miRFP670. To quantify cell morphology with high resolution, thin volume stacks of FUCCICardioids were acquired using the spinning disk confocal microscope at defined self-assembly stages. Datasets were denoised and deconvolved using the Richardson-Lucy algorithm (10 iterations) in Nikon NIS software. For each timepoint, a single actin focal plane at the basal region of the FUCCICardioids was selected. Actin boundaries were manually segmented using the Segmentation Editor in Fiji (ImageJ), and labeled masks were used to extract morphological parameters. Results are shown as mean ± standard error of the mean (SEM) in *Figure 2g*. Temporal changes in mean single-cell area ($\mu m^2$) were tracked from day 2.5 to late-stage (> day 25) differentiation. Statistical analysis revealed significant changes in cell area after day 2.5. All statistical analyses were performed using GraphPad Prism (v10.3.01) and SigmaPlot (v14.0), with results reported in the Data availability excel files. A nonparametric Kruskal-Wallis test was used to assess variation across timepoints, with Dunn's multiple comparison for post hoc analysis ($\alpha = 0.05$). Significant differences included Day 2.5 vs. Day 3.5: $p = 0.0004$; Day 2.5 vs. Day 4.5: $p < 0.0001$; Day 2.5 vs. > Day 25: $p = 0.0018$ (all comparisons shown in Figure 2g).

##### Cytotoxicity, microtubule dynamics, and nocodazole assays

The SPY650-tubulin probe (catalog# SC503, Spirochrome) was dissolved in 50 ul of dimethylsulphoxide (catalog# D2650-100ML, Merck) to obtain a 1000X stock solution. Nocodazole (catalog# M1404-50MG, Merck) was dissolved in dimethylsulphoxide at a concentration of 5 µg/ul. In the cytotoxicity and long-term cell proliferation experiments, hiPSC cardiomyocytes were loaded with SPY650-tubulin probe (1X) and incubated at 37 °C for 1h. Afterward, cardiomyocytes were imaged, calcium transients recorded (time 0, baseline) and subsequentially treated with Nocodazole at the final concentration of 5µM for 8h. The experiment was performed under live imaging conditions in an 8-well (catalog # 80807, Ibidi), and confocal imaging was performed using the Okolab enclosure to maintain a temperature of 37°C with 5% $CO_2$ and passive humidification. When 8h elapsed, cells were washed twice (washout) with PBS (without $CaCl_2$/$MgCl_2$) without altering the field of view, imaged, and calcium transients recorded (time 8h after dosing). The experiment was extended for an extra 12h and as for the previous time points, both structure and calcium were imaged at the end (20h) of the experiment. For cytotoxicity screening with hiPS-derived ectoderm, hiPSCs were plated in an 8-well ibidi slide coated with1:100 Growth Factor Reduced (GFR) Matrigel and differentiated for 7 days using the STEMdiff Trilineage Differentiation Kit (catalog# 05230, STEMCELL) accordingly with the manufacturer instructions. At the end of the differentiation, cells were treated with Nocodazole (100 nM) and imaged every 30 minutes for 18h. See Fig. S3 here and the relevant main and extended figures in the manuscript.

##### Microcontact printing of fibronectin on PDMS-coated dishes and coverslips

To confine single or few CMs in restricted and shape-controlled areas, we produced selective coatings of human fibronectin on polydimethylsiloxane (PDMS) substrate by microcontact printing[25]. As cell culture substrates, we first used off-the-shelf 35-mm-diameter dishes featuring a 100-µm-thick glass bottom coated with a 40-µm-thick layer of soft (1.5 kPa) PDMS (product number 81291, Ibidi). We also coated 25-mm-diameter round coverslips (thickness nr. 1, product number 06J0026, Knittel Glass), then coated with Sylgard 184 PDMS (Dow Corning Midland, MI). In detail, the coverslips were first cleaned with water and soap, then incubated in isopropanol (product number 8187661000, Sigma) for 60 s, and finally rinsed thoroughly in deionized water and dried. The base and curing agent of the commercial PDMS kit were mixed in 10:1 (w:w) and degassed for 1 h at <30 mbar. 1 mL of PDMS pre-polymer solution was cast on each coverslip and spin-coated as follows: 1000 rpm for 10 s, 2000 rpm for 10 s, and 3000 rpm for 30 s. After spin coating, the coverslips were placed on a hot plate at 120 °C for 1 h to cure and then stored inside a petridish to prevent contamination for further use[26].

The stamps for microcontact printing were fabricated using standard soft lithography techniques and Sylgard 184 PDMS, as previously described[27]. In particular, pillars with a rectangular base of 75 µm x

15 µm were used to produce islands of fibronectin on the otherwise adhesion-blocking PDMS surface. For sterilization before printing and cleaning after use, the PDMS stamps were immersed in a 70% ethanol (product number 93019.46.00, VWR) aqueous solution and sonicated for 15 minutes using a Branson system (Bransonic® M Mechanical Bath 1800).

For fibronectin loading, the PDMS stamps were removed from the solution and let dry inside a cell culture hood for at least 30 minutes. Fibronectin (either product number 341635, Sigma Aldrich, or product number 11051407001, Sigma Aldrich) was diluted to a working solution of 40 µg/mL and cast on the microstructured surface of the stamp for incubation (1 hour at room temperature in a sterile biological hood). After the incubation, the fibronectin solution was removed from the stamps, which were let dry for an hour. The recovered fibronectin solution was reused for the same process for a maximum of four printing iterations.

Before stamping, the PDMS surface of both commercial PDMS-coated dishes and prepared coverslips was treated using a UVO system (PSD-UV, Novascan) for 15 min. After the substrates were activated using UVO, the fibronectin-loaded stamps were inverted and gently placed on the PDMS surface for a few seconds. The microcontact-printed substrates were then left in the incubator overnight before cell seeding to allow the uncoated PDMS surface to recover its hydrophobic, cell-repellent properties. CMs were then seeded on the microcontact-printed substrates at a density (around 1 x $10^5$ cells per well) characterized to result in one cell per pattern on average. Imaging was performed 6-8 days after seeding as described in the previous sections.

**Micromolding of gelatin gels and formation of CM-aligned tissue**

To generate aligned CM tissues, we seeded CMs at high density on micromolded gelatin gels. PMDS molds were fabricated using the same process as described in the previous section, but with a different microstructure. Instead of pillars with a small island-like base, a linear array with 20 µm wide lines and 20 µm spacing between neighboring lines was selected. In terms of hydrogel composition, porcine gelatin (PG, type A, G2625, Sigma) was dissolved into Dulbecco's phosphate-buffered saline (PBS) without $Ca^{2+}$ and $Mg^{2+}$ (referred to as "PBS -/-," C-40232, PromoCell). The solution was heated to 50 °C and magnetically stirred (between 700 and 800 rpm) for at least 30 min using a hotplate stirrer until gelatin was fully dissolved. To enzymatically crosslink gelatin, Transglutaminase (TG) powder (Activa® TI Transglutaminase, 1002, Modernist Pantry) was dissolved in PBS -/-, vortexed (between 700 and 800 rpm) for at least 10 minutes until fully dissolved, and then filtered through a 0.45 µm filter. The PG and TG solutions were warmed to 37 °C, and mixed to reach a final concentration of 7.5% and 2% w/v, respectively. 35 mm diameter TC-treated COC-bottom µ-Dish (thickness 1.5, 81156, Ibidi) were used as substrate in combination with 50-µm-thick laser-cut steel (M-Tech®F, Georg Martin) spacers to control the hydrogel thickness. 200 µL hydrogel precursor solution was cast on the COC substrate, and the PDMS mold was inverted, placed on the PDMS surface, and at least 25 g weight was placed on each mold. The gel was left to set at room temperature overnight in a humidified environment (inside a petri dish with wet tissues). The following day, the molds were removed, and the resulting molded hydrogels were rinsed three times with deionized water to remove the enzyme and then incubated in CM media for 1 h before cell seeding with a density of 1 M cells per well. Imaging was performed 20 days after seeding as described in the previous sections.

**Analysis of calcium wave propagation on aligned CM tissues on molded gelatin gels**

20 days after seeding, the CMs seeded at high density (1 x $10^6$ cells/well) on gelatin hydrogels with molded line arrays were imaged using widefield mode using 10x objective (Plan Fluor 10x DIC L, Nikon) at 500 fps. Fields of view (FOVs) of roughly 1.4 mm x 1.4 mm were analyzed using a similar approach to what was previously described[28]. In essence, after flatfield and bleach correction the FOVs were processed to extract the activation time for each segmented area, construct chronomaps, and estimate the average propagation velocity.
